# GTPBP1 resolves paused ribosomes to maintain neuronal homeostasis

**DOI:** 10.1101/2020.09.03.281675

**Authors:** Markus Terrey, Scott I. Adamson, Alana L. Gibson, Tianda Deng, Ryuta Ishimura, Jeffrey H. Chuang, Susan L. Ackerman

**Affiliations:** Howard Hughes Medical Institute, Department of Cellular and Molecular Medicine, Section of Neurobiology, University of California San Diego, La Jolla, CA 92093, USA; Graduate School of Biomedical Sciences and Engineering, University of Maine, Orono, ME 04469, USA; The Jackson Laboratory for Genomic Medicine, Farmington, CT 06032, USA; Department of Genetics and Genome Sciences, Institute for Systems Genomics, UConn Health, Farmington, CT 06030, USA; Division of Biological Sciences Graduate Program, University of California San Diego, La Jolla, CA 92093, USA; The Jackson Laboratory, Bar Harbor, ME 04609, USA

## Abstract

Ribosome-associated quality control pathways respond to defects in translational elongation to recycle arrested ribosomes and degrade aberrant polypeptides and mRNAs. Loss of an individual tRNA gene leads to ribosomal pausing that is resolved by the translational GTPase GTPBP2, and in its absence causes neuron death. Here we show that loss of the homologous protein GTPBP1 during tRNA deficiency in the mouse brain also leads to codon-specific ribosome pausing and neurodegeneration, suggesting that these non-redundant translational GTPases function in the same pathway to mitigate ribosome pausing. Ribosome stalling in the mutant brain led to activation of the integrated stress response (ISR) mediated by GCN2 and decreased mTORC1 signaling. However, in contrast to the ISR, which enhanced neuron survival, reduced mTORC1 signaling increased neuronal death. Our data demonstrate that GTPBP1 functions as an important quality control mechanism during translation elongation and suggest that translational signaling pathways intricately interact to regulate neuronal homeostasis during defective translation elongation.

## Introduction

Translation of mRNA into protein utilizes four principle cycles: translation initiation, elongation, termination and ribosome recycling (Schuller and Green, 2019). During translation initiation, eukaryotic translation initiation factors (eIFs) mediate assembly of an 80S ribosome and an initiator methionyl-tRNA at the mRNA start codon. Subsequently, the elongating 80S ribosome moves along the mRNA decoding mRNA triplets until the mRNA termination codon is reached where eukaryotic peptide chain release factors (eRFs) stimulate the release of the nascent protein. Lastly, to permit engagement of ribosomes in new rounds of translation, the terminated 80S ribosome complex is separated into the 40S and 60S subunits and released from the mRNA by the ATP-binding cassette sub-family E member 1 (ABCE1).

Although much attention has been dedicated to the regulation of protein synthesis through translation initiation, collective evidence highlights the effect of translation elongation and its kinetics on protein synthesis (Stein and Frydman, 2019). Elongation rates not only vary between mRNAs generated from different genes but rates also fluctuate across a given transcript (Chaney and Clark, 2015; Kaiser and Liu, 2018; Rodnina, 2016). Multiple factors including secondary structure of mRNAs, availability of tRNAs, interactions of the nascent peptide with the ribosome and codon identity influence elongation rates, and variations in these rates affect co-translational protein folding, translational fidelity, and gene expression through mRNA decay (Brule and Grayhack, 2017; Buhr et al., 2016; Chaney and Clark, 2015; Drummond and Wilke, 2008; Rodnina, 2016; Spencer et al., 2012; Thommen et al., 2017; Wolf and Grayhack, 2015; Yu et al., 2015).

With growing evidence suggesting translation elongation is regulated, elaborate mRNA surveillance mechanism that resolve translation elongation defects have been identified (Brandman and Hegde, 2016; Joazeiro, 2019). We previously reported that an ENU-induced null mutation (*nmf205*) in the translational GTPase (trGTPase) GTPBP2 causes early, widespread neurodegeneration when present on the common inbred mouse strain C57BL/6J (B6J) (Ishimura et al., 2014). Neuron death in B6J-*Gtpbp2^nmf205/nmf205^* (B6J-*Gtpbp2*^-/-^) mice is a result of the epistatic interactions between the loss-of-function mutation in *Gtpbp2* and a hypomorphic mutation present in the B6J strain that disrupts processing of the brain-specific, nuclear encoded tRNA, *n-Tr20. n-Tr20* is widely expressed in neurons and is the most highly expressed member of the five tRNA^Arg^_UCU_ genes in brain, and the B6J-associated mutation reduces the pool of available tRNA^Arg^_UCU_ (Kapur et al., 2020). In the absence of *Gtpbp2*, translating ribosomes in the cerebellum exhibit prolonged pauses at AGA codons, suggesting GTPBP2 acts as a ribosome rescue factor to resolve codon-specific ribosome pausing occurring when the available pool of cognate tRNAs is limited. The essential role of *Gtpbp2* in neuronal homeostasis was further revealed by the identification of mutations in *Gtpbp2* causing neurological defects and intellectual disabilities in humans (Bertoli-Avella et al., 2018; Carter et al., 2019; Jaberi et al., 2016).

Defects in translation elongation may feedback to regulate translation initiation, supporting the emerging link between translation elongation and initiation to control global translation (Chu et al., 2014; Inglis et al., 2019; Liakath-Ali et al., 2018; Sanchez et al., 2019). Translation is highly regulated by two signaling pathways: the mTOR signaling pathway and the integrated stress response (ISR). mTORC1 phosphorylates the ribosomal protein S6 kinase (S6K1) and the eukaryotic translation initiation factor 4E binding protein 1 (4E-BP1) to positively regulate translation initiation and elongation (Showkat et al., 2014). The ISR reprograms the translatome by inhibiting translation initiation to suppress global translation via phosphorylation of the translation initiation factor eIF2α (p-eIF2α^S51^) which inhibits formation of the ternary complex, while allowing translation of stress-responsive genes such as ATF4 (Harding et al., 2003; Wortel et al., 2017). In the B6J-*Gtpbp2*^-/-^ cerebellum, ribosome stalling is accompanied by induction of the integrated stress response (ISR). Phosphorylation of eIF2α in the B6J-*Gtpbp2*^-/-^ cerebellum is mediated by GCN2, one of four known kinases in mammals which are activated during distinct cellular or environmental stressors. Deletion of *Gcn2* in B6J-*Gtpbp2*^-/-^ led to increased neurodegeneration, demonstrating GCN2-dependent activation of the ISR acts to partially restore cellular homeostasis (Chesnokova et al., 2017; Dalton et al., 2012; Ishimura et al., 2016).

In addition to *Gtpbp2*, the genome of many eukaryotes contains a related gene, *Gtpbp1* (Atkinson, 2015). A previous report demonstrated that loss of *Gtpbp1* in mice did not lead to overt defects, suggesting functional redundancy between *Gtpbp1* and *Gtpbp2* (Senju et al., 2000). However, here we demonstrate that loss of *Gtpbp1* in mice with the B6J-associated *n-Tr20* mutation have neurodegeneration identical to that observed in B6J-*Gtpbp2*^-/-^ mice. Furthermore, ribosome footprint profiling analysis suggests GTPBP1 functions as a novel ribosome rescue factor to resolve ribosome pausing defects during tRNA deficiency. As observed in B6J-*Gtpbp2*^-/-^ mice, the ISR is also induced in the *Gtpbp1*^-/-^ brain and protects neurons from ribosome-pausing induced neurodegeneration. Finally, we show that deficiencies in ribosome pause resolution alter mTOR signaling in a cell type-specific manner, suggesting that differences in the modulation of translational signaling pathways may contribute to the selective neurodegeneration observed with defects in translation elongation.

## Results

### Loss of *Gtpbp1* leads to widespread neurodegeneration when tRNA is deficient

Previously we demonstrated that loss of the trGTPase GTPBP2 causes progressive neurodegeneration in mice (Ishimura et al., 2014). However, mice homozygous for a null mutation in *Gtpbp1*, which encodes a structurally related trGTPase (*Figure 1A*), were reported to have no apparent phenotypes on a mixed genetic background (Senju et al., 2000). Both GTPBP2 and GTPBP1 are expressed in many tissues and *in situ* hybridization revealed that transcripts of these genes were widely expressed throughout the brain (Ishimura et al., 2014; Senju et al., 2000) (*Figure 1B-D, Figure 1 – figure supplement 1A - B*). Interestingly, expression of *Gtpbp1* and *Gtpbp2* occurred both in neurons that degenerate in *Gtpbp2*^-/-^ mice (e.g., cerebellar granule cells, dentate gyrus granule cells, neurons in cortical layer IV) and those that do not (e.g., hippocampal pyramidal cells and non-layer IV cortical neurons) suggesting possible functional redundancy between these genes in some cell types.

**Figure 1.**
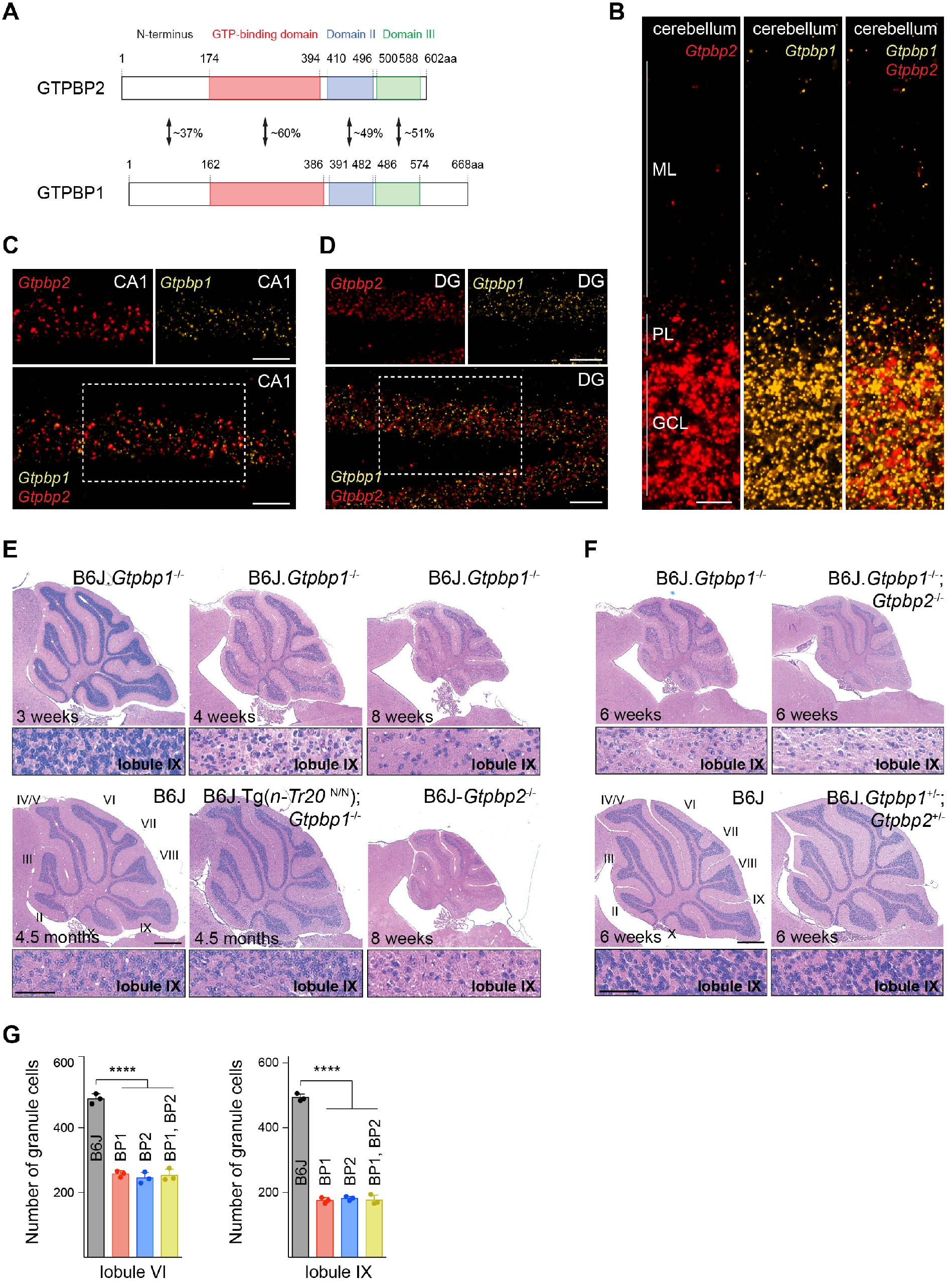
tRNA deficiency induces neurodegeneration in B6J.*Gtpbp1*^-/-^ mice. (*A*) Domain structure of mouse GTPBP2 and GTPBP1. The percent of identical amino acids between proteins are shown. (*B-D*) In situ hybridization demonstrating ubiquitous expression of *Gtpbp1* (yellow) and *Gtpbp2* (red) in the wild type (B6J) cerebellum (*B*), and areas of the hippocampus CA1 (*C*), dentate gyrus (DG) (*D*). Individual images of the areas defined by rectangles in *C* and *D* are shown above merged images. (*E, F*) Hematoxylin and eosin staining of sagittal sections of the cerebellum. Higher magnification images of lobule IX are shown below each genotype. Cerebellar lobules are indicated by Roman numerals. (*G*) Number of cerebellar granule cells of lobule VI and lobule IX at 6-week-old BP1 (B6J.*Gtpbp1*^-/-^); BP2 (B6J-*Gtpbp2*^-/-^); BP1, BP2 (B6J.*Gtpbp1*^-/-^; *Gtpbp2*^-/-^) mice. Data represent mean ± SD. ML, molecular cell layer; PL, Purkinje cell layer; GCL, granule cell layer. Scale bars: 20μm (*B, C, D*); 500μm and 50μm (higher magnification) (*E, F*). One-way ANOVA corrected for multiple comparisons using Tukey method (*G*). **** *P* ≤ 0.0001.

To investigate if phenotypes in *Gtpbp1*^-/-^ mice are dependent on genetic background as observed for B6J-*Gtpbp2*^-/-^ mice, we generated congenic B6J.*Gtpbp1*^-/-^ mice. Like B6J-*Gtpbp2*^-/-^ mice, B6J.*Gtpbp1*^-/-^ mice were indistinguishable from littermate controls at 3 weeks of age but developed overt ataxia by 6 weeks, and died by 8 weeks of age. Cerebellar degeneration was also similar to that observed in B6J-*Gtpbp2*^-/-^ mice (Ishimura et al., 2014) (*Figure 1E*). Apoptotic granule cells were observed in caudal lobes of the cerebellum just prior to 4 weeks of age and these neurons progressively died in a posterior to anterior manner. As previously observed in B6J-*Gtpbp2*^-/-^ mice, granule cells in the dentate gyrus (DG), layer IV cortical neurons and multiple neurons in the retina degenerated in B6J.*Gtpbp1*^-/-^ mice, but cell death was not observed for other neurons such as hippocampal pyramidal cells or neurons in the other layers of the cortex (*Figure 1 - figure supplement 2A-E*). Consistent with the dependency of cerebellar granule cell degeneration on levels of *n-Tr20*, a tRNA^Arg^_UCU_ gene with a processing mutation in the B6J strain, transgenic expression of wild type *n-Tr20*^N/N^ greatly attenuated degeneration of these neurons as previously observed for B6J-*Gtpbp2*^-/-^ mice (*Figure 1E*).

To genetically test for compensation between *Gtpbp1* and *Gtpbp2*, we analyzed mice that had mutations in both genes. Neurodegeneration was not observed in mice heterozygous for mutations in both *Gtpbp1* and *Gtpbp2* (*Figure 1F*). Furthermore, the onset, progression, and specificity of neurodegeneration were similar between B6J.*Gtpbp1*^-/-^; *Gtpbp2*^-/-^, B6J.*Gtpbp1*^-/-^, and B6J-*Gtpbp2*^-/-^ mice confirming that these genes are functionally distinct and do not compensate for each other in mediating cellular homeostasis that is necessary for neuron survival (*Figure 1F - G*).

**Figure 1 - figure supplement 1.**
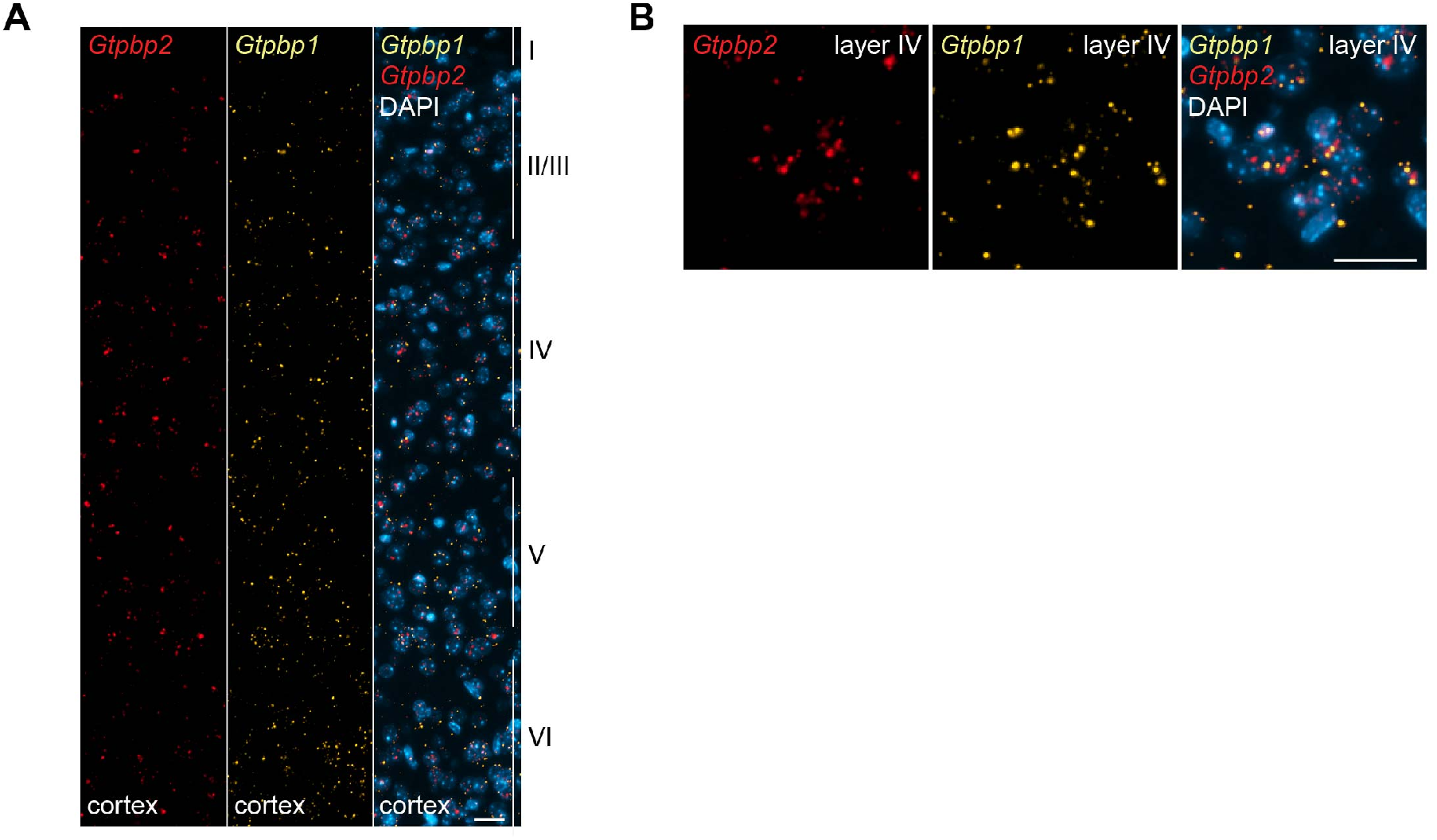
Expression of *Gtpbp1* and *Gtpbp2* in the mouse cortex. (*A* and *B*) In situ hybridization was performed on P28 B6J mice demonstrating ubiquitous expression of *Gtpbp1* (yellow) and *Gtpbp2* (red) throughout the cortex (*A*) and specifically in layer IV (*B*). Sections were counterstained with DAPI (blue). Scale bar: 20μm (*A, B*).

**Figure 1 - figure supplement 2.**
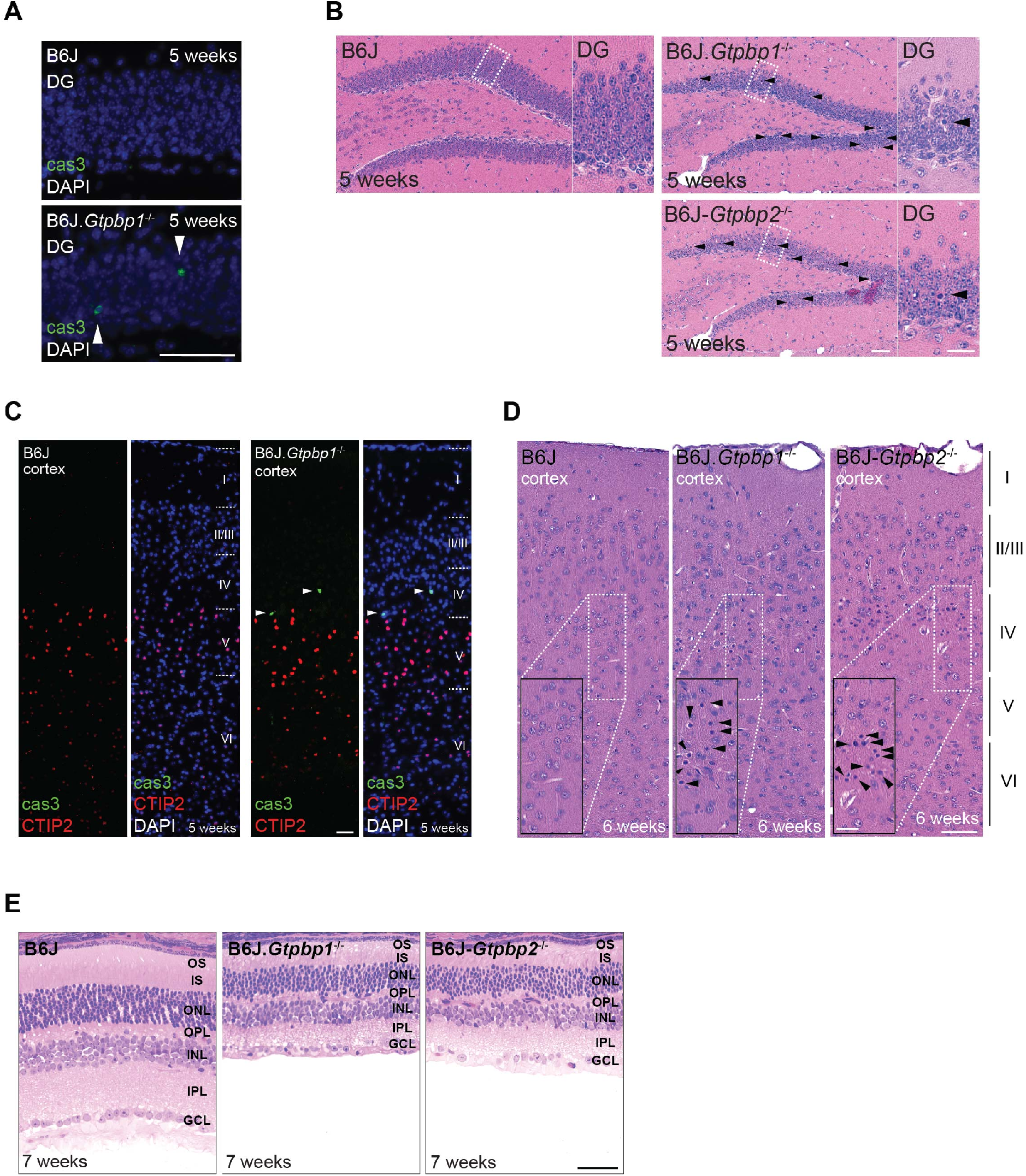
Extensive neurodegeneration in B6J.*Gtpbp1*^-/-^ mice. (*A*) Immunofluorescence of hippocampal sections of the dentate gyrus (DG) with antibodies against cleaved caspase 3 (cas3, green; arrowheads indicate positive cells). Sections were counterstained with DAPI. (*B*) Hematoxylin and eosin-stained sagittal sections of the dentate gyrus (DG). Higher magnification images of the region indicated by the rectangle are shown for each genotype. Arrowheads indicate pyknotic cells. Note the similarity in neurodegeneration between mutant genotypes. (*C*) Immunofluorescence on cortical sections with antibodies against cleaved caspase 3 (cas3, green) and CTIP2 (red) to identify layer V neurons. Sections were counterstained with DAPI. Cortical layers are indicated by Roman numerals. Note the cas3-positive neurons in layer IV (above layer V CTIP2^+^ neurons). (*D*) Hematoxylin and eosin-stained sagittal sections of the cortex from 6-week-old mice. Cortical layers are indicated by Roman numerals and higher magnifications (dashed line rectangle) of layer IV of the cortex are shown. (*E*) Hematoxylin and eosin-stained retina sections. Note the similarity in neurodegeneration between mutant genotypes. Scale bars: 50μm (*A, E*); 25μm (*C*); 50μm and 20μm (higher magnification) (*B, D*). OS, outer segment; IS, inner segment; ONL, outer nuclear layer; OPL, outer plexiform layer; INL, inner nuclear layer; IPL, inner plexiform layer; GCL, ganglion cell layer.

### GTPBP1 is a novel ribosome rescue factor

The genetic interaction between the B6J-derived *n-Tr20* mutation and the loss of *Gtpbp1* suggested that like GTPBP2, GTPBP1 might act as a rescue factor for ribosomes paused at AGA codons. To investigate this possibility, we performed ribosome profiling on cerebella from 3-week-old B6J.*Gtpbp1*^-/-^ and B6J mice. As previously observed in the B6J-*Gtpbp2*^-/-^ cerebellum, ribosome occupancy dramatically increased in the B6J-*Gtpbp1*^-/-^ cerebellum when AGA codons were in the A site of the ribosome (*Figure 2A - B, Figure 2 - figure supplement 1A*). Comparison of genes with AGA pauses revealed that approximately 50% of pausing genes were shared between libraries generated from individual B6J.*Gtpbp1*^-/-^ mice, and this was also true for B6J-*Gtpbp2*^-/-^ mice, supporting that AGA pausing is likely stochastic (*Figure 2C*). Similarly, about 50% of AGA pausing genes with increased AGA occupancy intersected between B6J.*Gtpbp1*^-/-^ and B6J-*Gtpbp2*^-/-^ mice (*Figure 2 - figure supplement 1B, Table S1*). Gene ontology (GO) analysis of pausing genes revealed enrichment in numerous biological processes and the majority of these processes were enriched in both *Gtpbp1* and *Gtpbp2* (*Figure 2 - figure supplement 1C*). Thus, our data suggest that both GTPBP1 and GTPBP2 rescue stochastic ribosome pausing events when the tRNA^Arg^_UCU_ pool is reduced.

**Figure 2.**
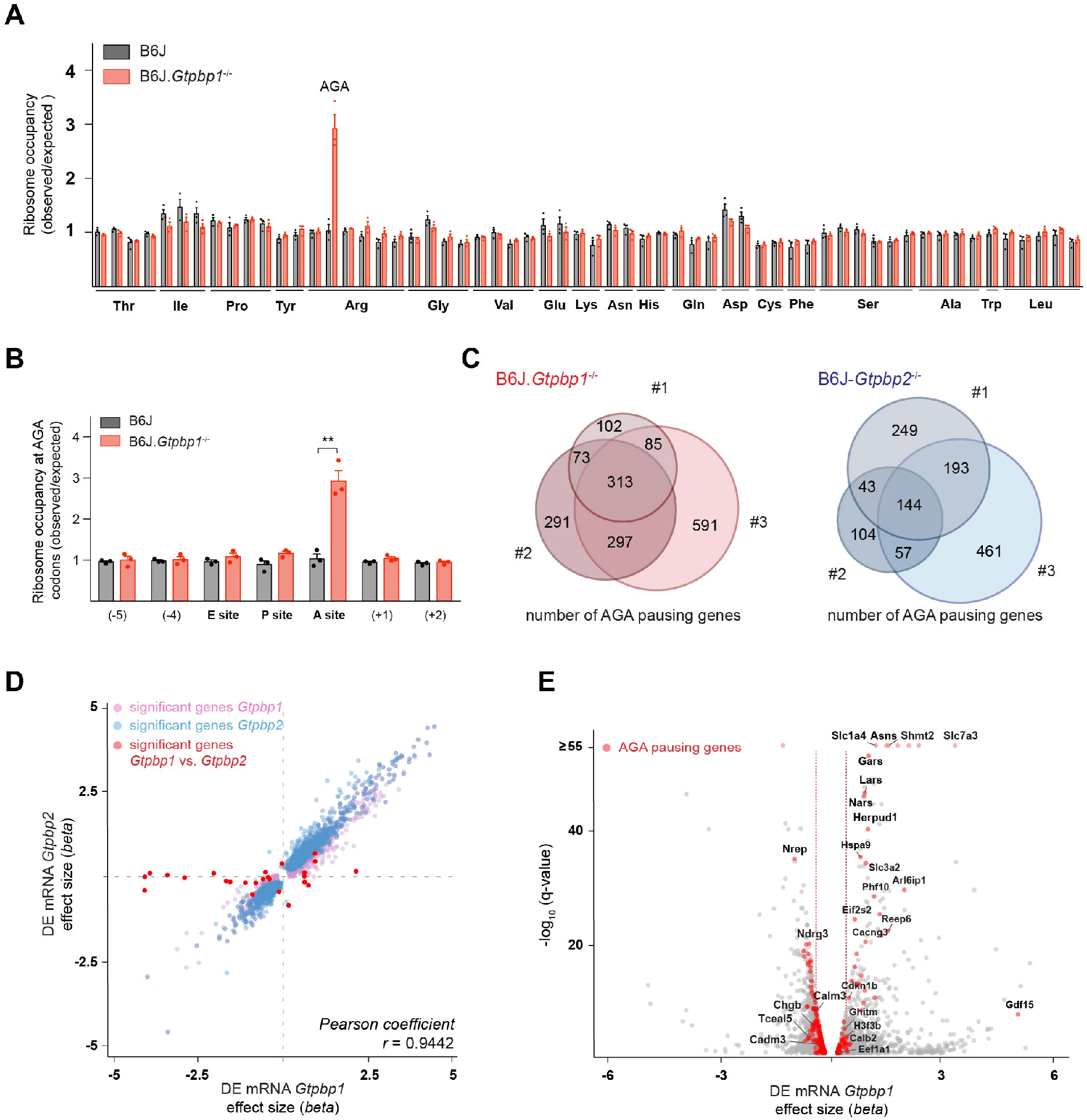
GTPBP1 resolves ribosome pausing induced by tRNA deficiency. (*A*) Ribosome occupancy was calculated by dividing the number of genome-wide reads at codons by the expected reads in the ribosomal A-site. Data represent mean ± SEM. Note that ribosome occupancy increased only at AGA codons in B6J.*Gtpbp1*^-/-^ mice. (*B*) Ribosome occupancy at AGA codons was calculated by dividing genome-wide reads at AGA codons by expected reads. Data represent mean ± SEM from ribosome profiling from cerebella of P21 B6J and B6J.*Gtpbp1*^-/-^ mice. (*C*) Venn diagram of genes with increased ribosome occupancy at AGA codons (z-score ≥10) between libraries prepared from individual B6J.*Gtpbp1*^-/-^ (red) or B6J-*Gtpbp2*^-/-^ (blue) mice (biological replicates 1-3). (*D*) Analysis of differential gene expression (DE mRNA) between B6J-*Gtpbp2*^-/-^ and B6J.*Gtpbp1*^-/-^ mice. Significant (q-value ≤ 0.05) gene expression changes are shown. *Gtpbp1*-dependent changes in pink; *Gtpbp2*-dependent changes in blue; and significant changes between *Gtpbp1* and *Gtpbp2* are shown in red (27 genes). The *beta* effect size is analogous to the natural log fold change in gene expression. (*E*) Analysis of differential gene expression (DE mRNA) between B6J and B6J.*Gtpbp1*^-/-^ mice. Significant (q-value ≤ 0.05) gene expression changes are shown in grey and genes with increased AGA ribosome occupancy (z-score ≥10, detected in at least 2 biological replicates) that are differentially expressed are highlighted in red (260 genes). The *beta* effect size is analogous to the natural log fold change in gene expression and 1.5-fold changes in gene expression are indicated by the red dashed lines. Multiple *t* tests corrected for multiple comparisons using Holm-Sidak method (*B*). ** *P* ≤ 0.01.

To determine if transcriptional alterations are shared upon loss of *Gtpbp1* and *Gtpbp2*, we performed RNA-Sequencing analysis (Table S2). Comparison of data from the cerebellum of 3-week-old B6J.*Gtpbp1*^-/-^ and B6J mice revealed differential expression (q-value ≤ 0.05) of approximately 12% of detected genes, with 46% upregulated and 54% downregulated. Similarly, loss of *Gtpbp2* altered expression of about 10% of detected genes with 45% and 55% of genes upregulated and downregulated, respectively. Interestingly, only 27 genes (mostly non-coding transcripts) were differentially expressed between *Gtpbp1*^-/-^ and *Gtpbp2*^-/-^ cerebella, revealing that loss of either gene similarly alters the transcriptome (*Figure 2D*).

The high similarity of gene expression changes in *Gtpbp1* and *Gtpbp2* mutant mice, combined with our observation that ribosome pausing defects are likely stochastic, suggested that transcript alterations might reflect a common cellular response to ribosome pausing rather than changes in levels of the specific mRNAs that harbor paused ribosomes. In agreement, gene expression changes were only weakly correlated with the transcripts showing increased ribosomal occupancy (Spearman correlation coefficient of 0.0405, p-value = 0.0083), with significant (q-value ≤ 0.05) changes in the transcript levels of only 33% or 26% of the genes with AGA pauses (that were detected in at least 2 replicates) in the *Gtpbp1*^-/-^ and *Gtpbp2*^-/-^ cerebellum, respectively (*Figure 2E*).

**Figure 2 - figure supplement 1.**
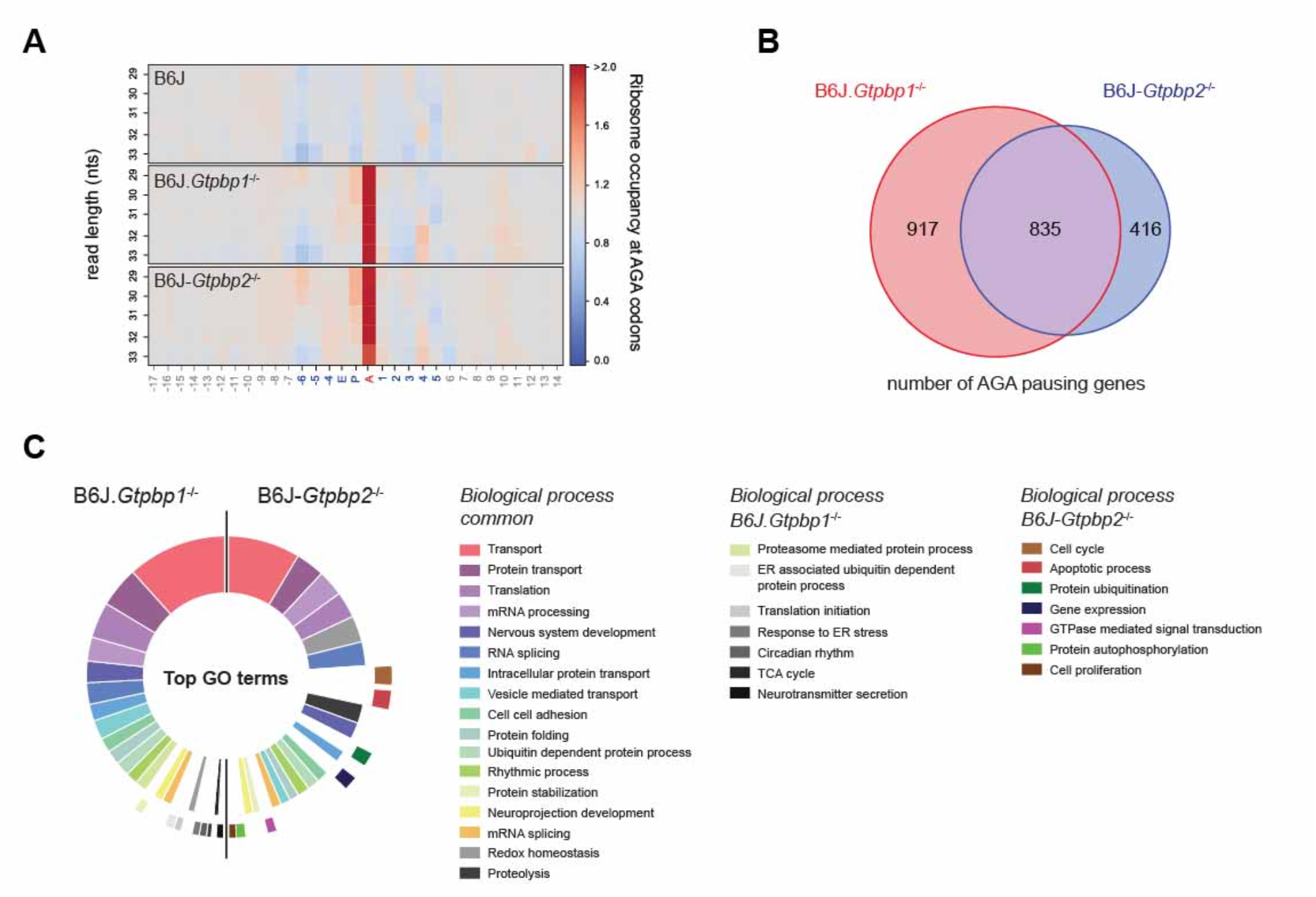
*Gtpbp1* and *Gtpbp2* resolve AGA pauses. (*A*) Heatmap of ribosome occupancy (observed / expected reads) at AGA codons relative to nucleotide (nts) length of ribosome protected fragments. Note the increase in ribosome occupancy at the A-site AGA codon in B6J.*Gtpbp1*^-/-^ and B6J-*Gtpbp2*^-/-^ mice. (*B*) Venn diagram of all genes with increased AGA ribosome occupancy (z-score ≥10) observed in B6J.*Gtpbp1*^-/-^ and B6J-*Gtpbp2*^-/-^ mice. (*C*) Gene Ontology (GO) analysis using all AGA pausing genes (z-score ≥10) from B6J.*Gtpbp1*^-/-^ and B6J-*Gtpbp2*^-/-^ mice. Top 24 enriched categories (Benjamini-Hochberg adjusted p-value ≤ 0.05) are shown. The inner circle and outer circle represent GO categories that are shared between mutant genotypes or unique to each genotype, respectively.

### Loss of GTPBP1 activates the integrated stress response

In order to identify molecular pathways that might respond to ribosome pausing, we performed upstream regulator analysis (Ingenuity Pathway Analysis) of genes differentially expressed between B6J and B6J.*Gtpbp1*^-/-^ mice. This analysis revealed significant enrichment for activation of the transcription factor ATF4, an effector of the integrated stress response (ISR) (*Figure 3A*, inset). The ISR is induced by the coupling of stress signals to the phosphorylation of serine 51 of the translation initiation factor eIF2α to decrease translation initiation. While the ISR reduces translation of many mRNAs, translation of ATF4 is enhanced, which results in the upregulation of genes to restore cellular homeostasis. We previously observed induction of the ISR in the B6J-*Gtpbp2*^-/-^ cerebellum suggesting that activation of this pathway may be a common cellular response to ribosome stalling (Ishimura et al., 2016). In agreement, *Atf4* and 141 of the 153 differentially expressed ATF4 target genes (Jaeseok Han et al., 2013) were upregulated in the cerebellum of 3-week-old B6J.*Gtpbp1*^-/-^ mice (*Figure 3A*). Furthermore, levels of phosphorylated eIF2α (p-eIF2α^S51^) were increased in the cerebellum and hippocampus of B6J.*Gtpbp1*^-/-^ mice (*Figure 3B - C*). In agreement, *in situ* hybridization of the B6J.*Gtpbp1*^-/-^ hippocampus with probes to induced ATF4 targets in the cerebellum of these mice demonstrated that these transcripts were also induced in hippocampal neurons of 3-week-old mutant mice (*Figure 3D*). Interestingly, induction of these genes varied in different types of neurons. *Sens2, Slc7a1*, and *Chac1* were upregulated in both CA1 and DG neurons, whereas *Ddr2* was only upregulated in CA1 neurons (*Figure 3D*).

**Figure 3.**
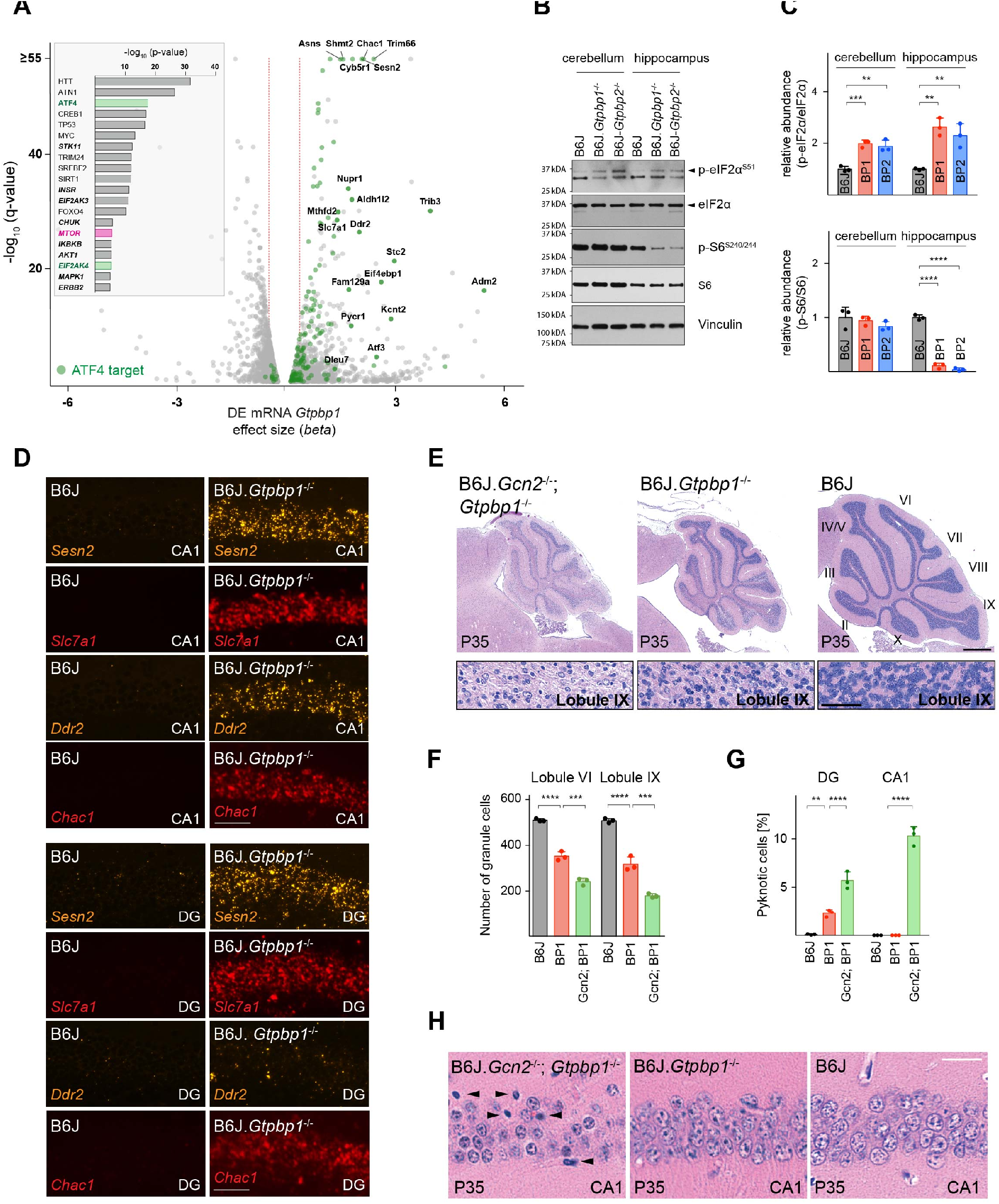
Ribosome pausing activates the integrated stress response (ISR) to ameliorate neurodegeneration in *Gtpbp1*^-/-^ mice. (*A*) Analysis of differential gene expression between B6J and congenic B6J.*Gtpbp1*^-/-^ mice. Significant (q-value ≤ 0.05) gene expression changes are shown in grey and differentially expressed ATF4 target genes are highlighted in green. The *beta* effect size is analogous to the natural log fold change in gene expression and 1.5-fold changes in gene expression are indicated (red dashed lines). (Inset) Identification of upstream regulators using Ingenuity Pathway Analysis (IPA) of differentially expressed genes between B6J and B6J.*Gtpbp1*^-/-^ mice. Top ten transcription factors and kinases (***italics***) are shown. (*B*) Western blotting analysis of tissue lysates from P21 mice. Vinculin was used as an input control. (*C*) Relative abundance of p-eIF2α^S51^ and p-S6^S240/244^ in the hippocampus and cerebellum of BP1 (B6J.*Gtpbp1*^-/-^), BP2 (B6J-*Gtpbp2*^-/-^), and control (B6J) mice. Levels of p-eIF2α^S51^ or p-S6^S240/244^ were normalized to total level of eIF2α or S6 and phosphorylation levels are relative to those of B6J. Data represent mean ± SD. (*D*) *In situ* hybridization of ATF4 targets of the hippocampal CA1 and dentate gyrus (DG). (*E*) Sagittal cerebella sections stained with hematoxylin and eosin. Higher magnification images of lobule IX are shown below each genotype. Cerebellar lobes are indicated by Roman numerals. (*F*) Quantification of cerebellar granule cells of lobule VI and lobule IX of control (B6J), BP1 (B6J.*Gtpbp1*^-/-^), and Gcn2; BP1 (B6J*.Gcn2*^-/-^; *Gtpbp1*^-/-^) mice at P35. Data represent mean ± SD. (*G*) Percent of dentate gyrus (DG) and CA1 neurons that are pyknotic in control (B6J), BP1 (B6J.*Gtpbp1*^-/-^), and Gcn2; BP1 (B6J*.Gcn2*^-/-^; *Gtpbp1*^-/-^) mice at P35. Data represent mean ± SD. (*H*) Hippocampal sagittal sections of the CA1 stained with hematoxylin and eosin. Arrowheads indicate pyknotic cells. Scale bars: 50μm (*D*); 500μm and 50μm (higher magnification) (*E*); 20μm (*H*). One-way ANOVA, corrected for multiple comparisons using Tukey method (*C, F, G*). ** *P* ≤ 0.01, *** *P* ≤ 0.001, **** *P* ≤ 0.0001.

In B6J-*Gtpbp2*^-/-^ mice the ISR is activated by the eIF2α kinase GCN2 (eIF2α K4). Inhibition of ISR activation via *Gcn2* deletion in these mice accelerated cerebellar granule cell death and induced neurodegeneration of hippocampal pyramidal cells (Ishimura et al., 2016). GCN2 was enriched in upstream regulator analysis of differentially expressed transcripts in the B6J.*Gtpbp1*^-/-^ cerebellum, suggesting that this kinase may also mediate activation of the ISR in *Gtpbp1*-deficient mice (*Figure 3A*, inset). In agreement, *Gcn2* deletion in B6J.*Gtpbp1*^-/-^ mice accelerated degeneration of cerebellar granule cells. In 5-week-old B6J.*Gtpbp1*^-/-^; *Gcn2*^-/-^ mice, 65% of these neurons had degenerated compared to 37% in B6J.*Gtpbp1*^-/-^ mice (*Figure 3E - F*). Furthermore, the dentate gyrus of B6J.*Gtpbp1*^-/-^; *Gcn2*^-/-^ mice had twice the number of granule cells with pyknotic nuclei compared to age-matched B6J.*Gtpbp1*^-/-^ mice (*Figure 3G, Figure 3 - figure supplement 1*). Finally, although CA1 pyramidal neurons do not normally degenerate in B6J.*Gtpbp1*^-/-^ mice, 10.3% of these neurons were undergoing apoptosis in the hippocampus of 5-week-old B6J.*Gtpbp1*^-/-^; *Gcn2*^-/-^ mice (*Figure 3G - H*). As previously reported (Ishimura et al., 2016; Zhang et al., 2002), neurodegeneration was not observed in the B6J.*Gcn2*^-/-^ brain (data not shown).

**Figure 3 - figure supplement 1.**
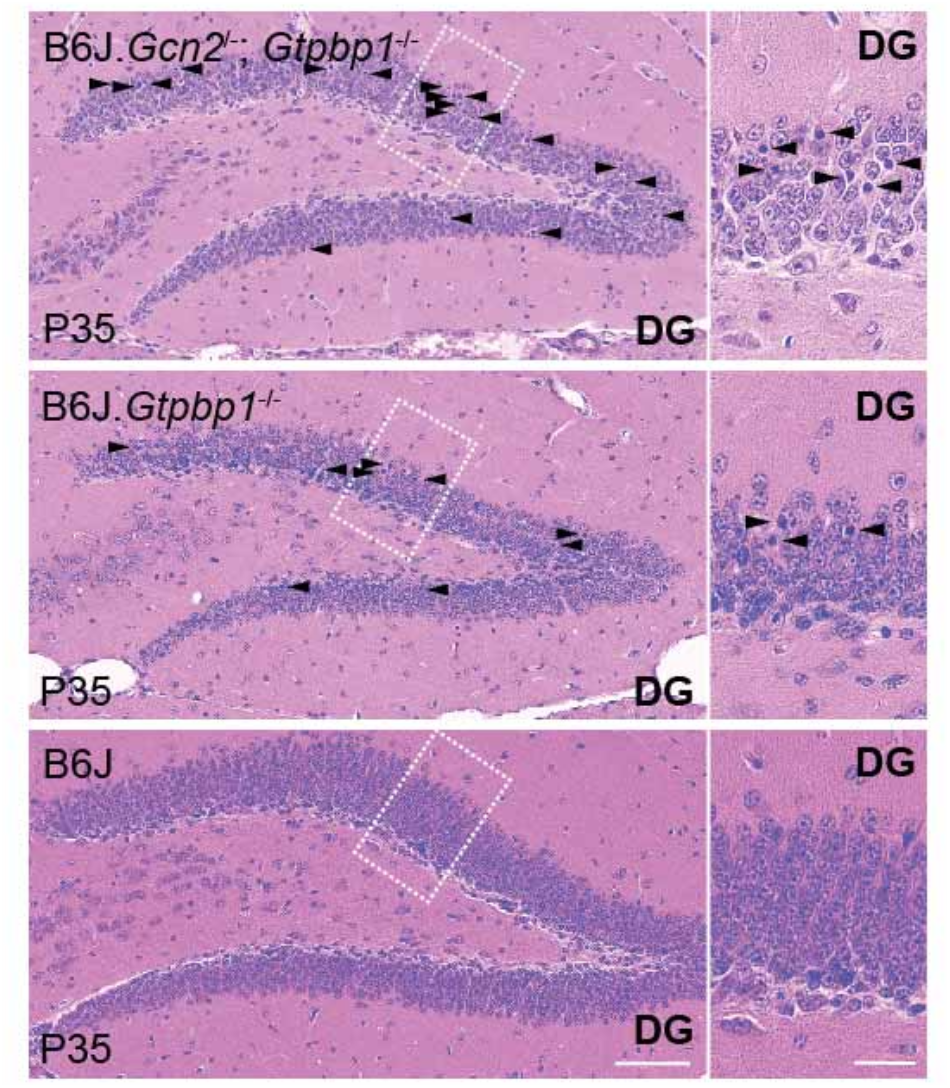
Loss of GCN2 enhances hippocampal degeneration in *Gtpbp1*^-/-^ mice. Hippocampal sagittal sections of the dentate gyrus (DG) stained with hematoxylin and eosin. Higher magnification images of the dentate gyrus region defined by the rectangle are shown. Arrowheads indicate pyknotic cells. Scale bars: 50μm and 20μm (higher magnification).

### Ribosome stalling induces neuron-specific changes in mTOR signaling

Our data suggest that defects in translation elongation activate the ISR, which regulates translation initiation and influences neurodegeneration. Thus, we wondered if ribosome pausing in *Gtpbp1* or *Gtpbp2* mutant mice could alter additional translational control pathways. Upstream regulator analysis of differentially expressed genes in the cerebellum of B6J.*Gtpbp1*^-/-^ mice revealed enrichment, although lower when compared to ATF4, of the mammalian target of rapamycin kinase (mTOR) (*Figure 3A*, inset). mTOR functions in two distinct protein complexes known as mTOR complex 1 (mTORC1) and complex 2 (mTORC2). Although association with membrane bound ribosomes (i.e., the endoplasmic reticulum) has been observed for mTORC2, the central role of mTOR in protein translation is largely attributed to mTORC1 through phosphorylation of specific effector proteins and translation of discrete mRNAs which contain 5’ terminal oligopyrimidine (TOP) tracts (Dai and Lu, 2009; Laplante and Sabatini, 2013, 2009; Ma and Blenis, 2009a, 2009b; Nandagopal and Roux, 2015; Zinzalla et al., 2011). To determine if mTOR activity is altered by loss of *Gtpbp1* or *Gtpbp2*, we analyzed the translational efficiency (TE) (Table S3) of mRNAs regulated by mTORC1 via their TOP motifs and assessed the phosphorylation status of the ribosomal protein S6 (p-S6^240/244^), a known downstream target of mTORC1 (Saxton and Sabatini, 2017; Thoreen et al., 2012). Only one of the 53 detected 5’TOP mRNAs (*Rps29*) had an altered TE in the B6J.*Gtpbp1*^-/-^ cerebellum and none of these genes displayed TE alterations in the B6J-*Gtpbp2*^-/-^ cerebellum (*Figure 4 - figure supplement 1A*). In addition, levels of p-S6^240/244^ were unchanged at 3 weeks of age in the cerebellum of both mutant strains, suggesting that mTOR signaling is not altered by ribosome pausing in granule cells, which comprise the vast majority of cells in the cerebellum (*Figure 3B - C*).

In contrast to the cerebellum, p-S6^240/244^ levels were decreased by about 90% in the hippocampus of 3-week-old B6J.*Gtpbp1*^-/-^ and B6J-*Gtpbp2*^-/-^ relative to wild type mice (*Figure 3B - C*). Interestingly, immunofluorescence using antibodies to p-S6^240/244^ revealed that the reduction of p-S6^240/244^ in the mutant hippocampus was more pronounced in specific neuron populations. In the control hippocampus, p-S6^240/244^ signal was most intense in CA3 pyramidal cells and lower in CA1 pyramidal cells and dentate gyrus granule cells (*Figure 4A*). In the mutant hippocampus, p-S6^240/244^ levels were dramatically reduced in the granule cells of the dentate gyrus and some scattered CA1 neurons (*Figure 4A*). In addition, levels of p-S6^240/244^ were not affected in CA3 or cortical neurons (i.e. layer IV cortical neurons) of 3-week-old B6J.*Gtpbp1*^-/-^ and B6J-*Gtpbp2*^-/-^ mice (*Figure 4 - figure supplement 1B - C*). Together, these results demonstrate that changes in mTOR signaling may occur in a cell-type specific manner in the brain of B6J.*Gtpbp1*^-/-^ and B6J-*Gtpbp2*^-/-^ mice.

**Figure 4.**
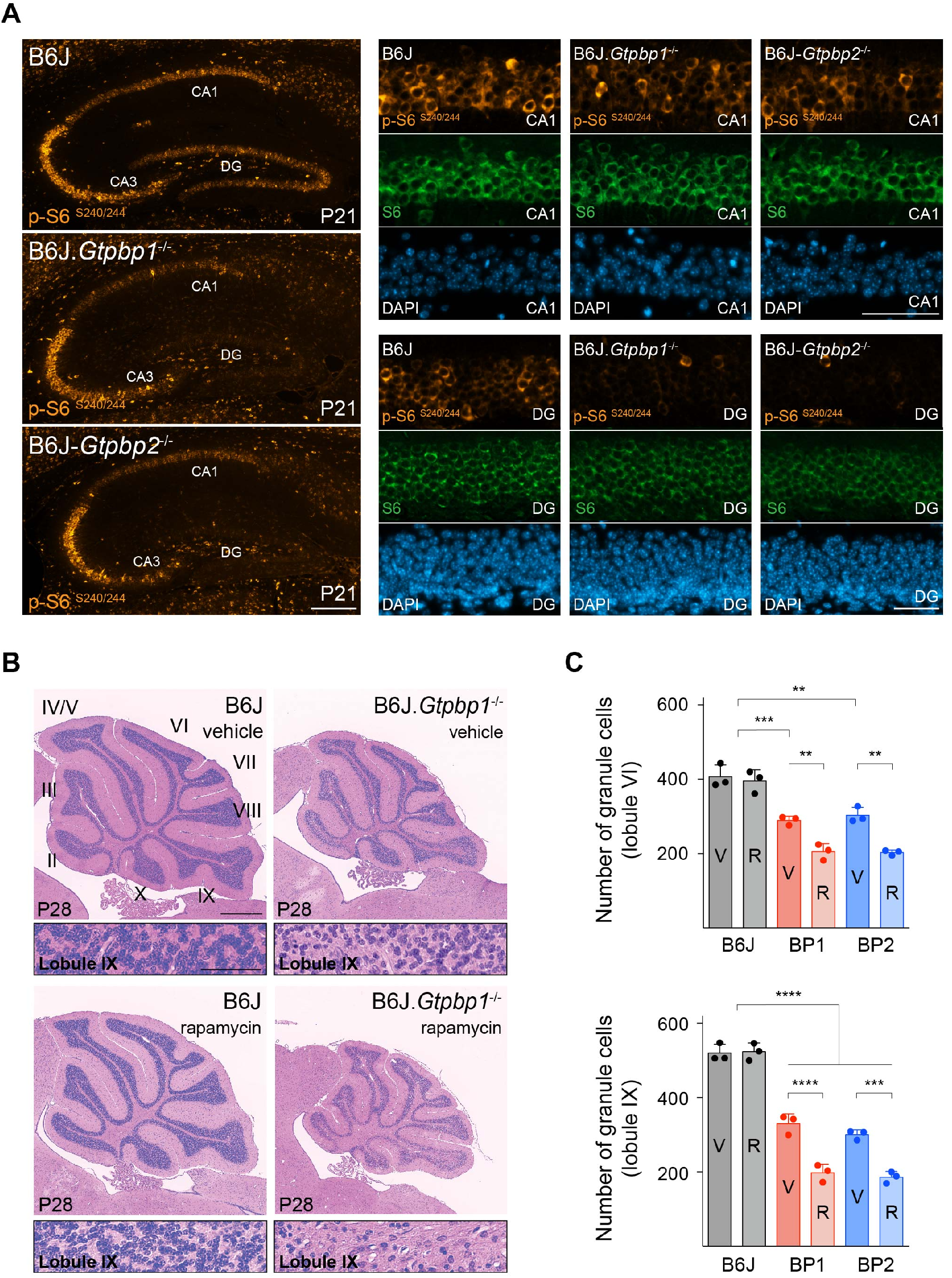
Decreased mTOR signaling enhances neurodegeneration in trGTPase-deficient mice. (*A*) Immunofluorescence of hippocampal sections with antibodies against p-S6^S240/244^ (orange), S6 (green). Sections were counterstained with DAPI (blue). Higher magnifications of CA1 and dentate gyrus (DG) are shown. (*B*) Sagittal cerebella sections of P28 mice injected with vehicle or rapamycin for 14 days stained with hematoxylin and eosin. Higher magnification images of lobule IX are shown below each genotype. Cerebellar lobes are indicated by Roman numerals. (*C*) Quantification of cerebellar granule cells in lobule VI and lobule IX of either vehicle (V) or rapamycin (R) treated control (B6J), BP1 (B6J.*Gtpbp1*^-/-^), and BP2 (B6J-*Gtpbp2*^-/-^) mice. Data represent mean ± SD. Scale bars: 100μm and 50μm (higher magnification) (*A*); 500μm and 50μm (higher magnification) (*B*). Two-way ANOVA corrected for multiple comparisons using Tukey method (*C*). ** *P* ≤ 0.01, *** *P* ≤ 0.001, **** *P* ≤ 0.0001.

The decrease in translation initiation by inhibition of the mTOR pathway has been observed to reduce ribosome pausing during amino acid deprivation (Darnell et al., 2018) suggesting that like the ISR, this pathway may also be protective in the *Gtpbp1* and *Gtpbp2* mutant brain. However, changes in levels of p-S6^240/244^ in the mutant hippocampus suggested that decreases in mTOR activity were most profound in hippocampal neurons that ultimately degenerate (i.e. the granule cells of the dentate gyrus). Thus, to determine the role of mTOR signaling on neuron survival during ribosome pausing, we pharmacologically inhibited mTOR by treating B6J, B6J.*Gtpbp1*^-/-^ and B6J-*Gtpbp2*^-/-^ mice with rapamycin daily for two weeks beginning at P14. Similar to the dramatic decrease in p-S6^240/244^ in 3-week-old hippocampi in B6J.*Gtpbp1*^-/-^ and B6J-*Gtpbp2*^-/-^ mice, cerebellar levels of p-S6^240/244^ were decreased by approximately 85% by P21 in rapamycin-treated mice (*Figure 4 - figure supplement 1D-F*). Examination of the cerebellum from P28 B6J.*Gtpbp1*^-/-^ and B6J-*Gtpbp2*^-/-^ cerebellum revealed that rapamycin treatment increased granule cell loss by 30% (lobule VI) and 40% (lobule IX) compared to mutant mice treated with vehicle (*Figure 4B - C*). No neuron loss was observed in rapamycin-treated control (B6J) mice (*Figure 4B - C*). Although it has been reported that mTOR inhibition may cause repression of *Atf4* transcripts and some of its target genes (Park et al., 2017), no significant reduction of *Atf4* or its targets was observed in rapamycin-treated mutant mice, suggesting that the ISR and mTOR pathways function independently upon loss of *Gtpbp1* and *Gtpbp2* in the B6J cerebellum (*Figure 4 - figure supplement 1G*).

Rapamycin treatment also increased apoptosis of layer IV cortical neurons by about 3-fold in P28 B6J.*Gtpbp1*^-/-^ and B6J-*Gtpbp2*^-/-^ mice but did not cause death of other cortical neurons (*Figure 4 - figure supplement 2A - B*). In addition, death of granule cells in the dentate gyrus was accelerated upon rapamycin treatment. Pyknotic nuclei were observed in 22% of these neurons in P28 rapamycin-treated B6J.*Gtpbp1*^-/-^ or B6J-*Gtpbp2*^-/-^ mice whereas only 3.4% of these neurons had pyknotic nuclei in vehicle-treated mutant mice (*Figure 4 - figure supplement 2C - D*). In addition, the number of granule cells was decreased in the dentate gyrus of rapamycin-treated mice by 35%, indicating apoptosis in this region of the brain began earlier in rapamycin-treated mutant mice (*Figure 4 - figure supplement 2E*). Finally, apoptosis of CA1 neurons was not observed in rapamycin-treated B6J.*Gtpbp1*^-/-^ and B6J-*Gtpbp2*^-/-^ mice at P28 when treatment began at P14. However, pyknotic nuclei were observed when mice were treated from P28-P42, suggesting that while these neurons are sensitive to mTOR suppression, the timing of this sensitivity differs from that of other neurons (*Figure 4 - figure supplement 2F*). Taken together, our data demonstrate that unlike the ISR, which acts to prevent loss of neurons with ribosome stalling, inhibition of mTOR increases the vulnerability of multiple neuronal populations.

**Figure 4 - figure supplement 1.**
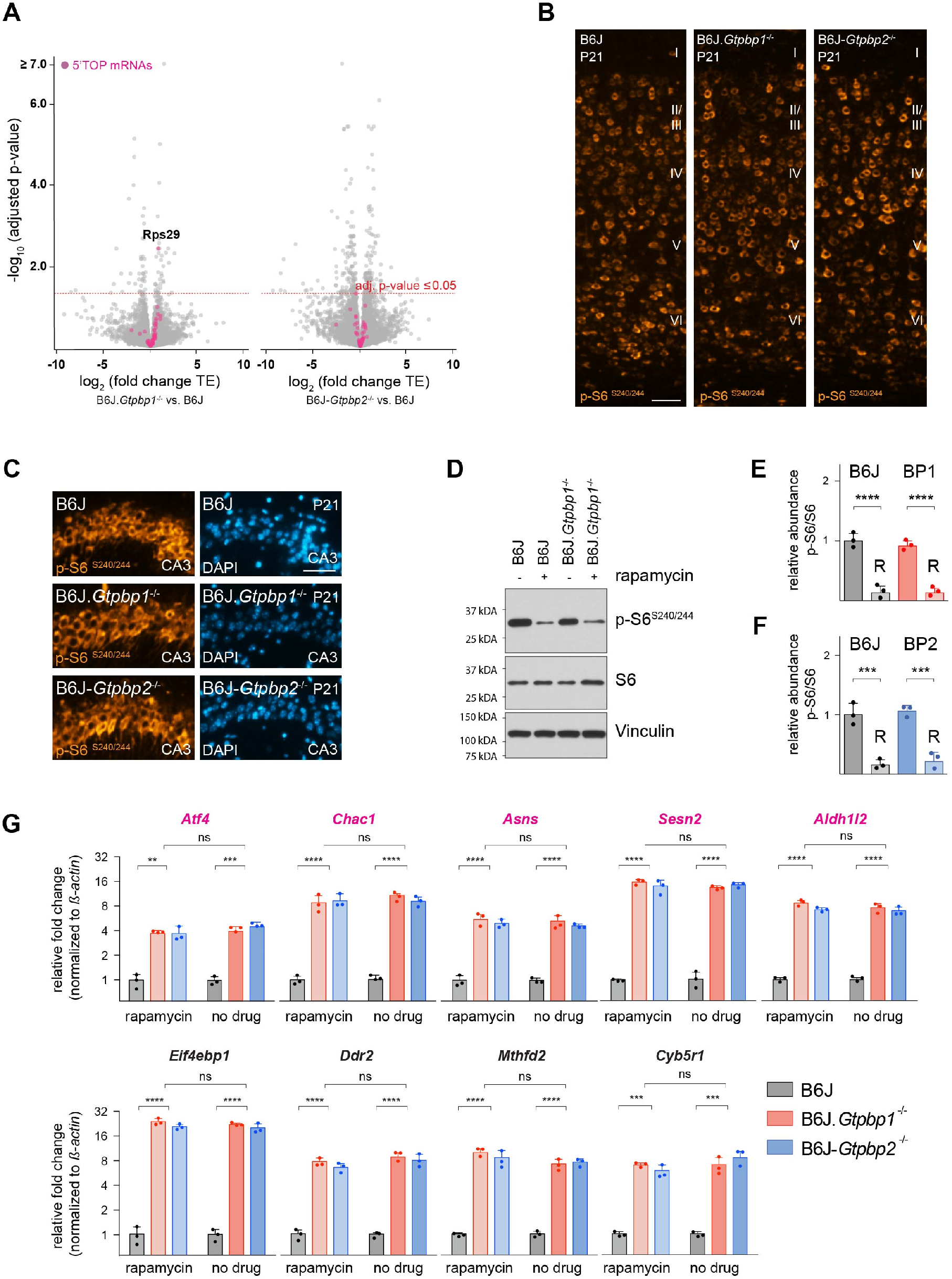
Analysis of mTOR signaling in trGTPase-deficient mice. (*A*) Analysis of differential translation efficiency (TE) in the cerebellum of B6J and B6J.*Gtpbp1*^-/-^ or B6J-*Gtpbp2*^-/-^ mice. 5’TOP mRNAs (53 genes) regulated by mTOR are shown in magenta. The red dashed line marks significant changes in TE (adjusted p-value ≤ 0.05). (*B*) Immunofluorescence of cortical sections with antibodies against p-S6^S240/244^ (orange). (*C*) Immunofluorescence of hippocampal (CA3) sections with antibodies against p-S6^S240/244^ (orange) and counterstained with DAPI (blue). (*D*) Western blotting analysis of p-S6^S240/244^ in cerebellar tissue extracts from P21 B6J and B6J.*Gtpbp1*^-/-^ mice untreated or injected with rapamycin for 7 days. Vinculin was used as an input control. (*E*) Relative abundance of p-S6^S240/244^ in the cerebellum of B6J and BP1 (B6J.*Gtpbp1*^-/-^) mice treated with rapamycin (R). Levels of p-S6^S240/244^ were normalized to total level of S6 and phosphorylation levels are relative to that of untreated B6J mice. Data represent mean ± SD. (*F*) Relative abundance of p-S6^S240/244^ in the cerebellum of B6J and BP2 (B6J-*Gtpbp2*^-/-^) mice treated with rapamycin (R). Levels of p-S6^S240/244^ were normalized to total level of S6 and phosphorylation levels are relative to that of untreated B6J mice (unmarked bar). Data represent mean ± SD. (*G*) Quantitative RT-PCR analysis of ATF4 targets using cerebellar cDNA from P21 B6J, B6J.*Gtpbp1*^-/-^, and B6J-*Gtpbp2*^-/-^ mice injected with rapamycin for 7 days or untreated. Data were normalized to beta-actin and fold change in gene expression is relative to that of B6J. Magenta colored genes were previously shown to be sensitive to mTOR inhibition (Park et al., 2017). Data represent mean ± SD. Scale bar: 50μm (*B, C*). Twoway ANOVA (*E, F, G*), corrected for multiple comparisons using Tukey method. ns, not significant, ** *P* ≤ 0.01, *** *P* ≤ 0.001, **** *P* ≤ 0.0001.

**Figure 4 - figure supplement 2.**
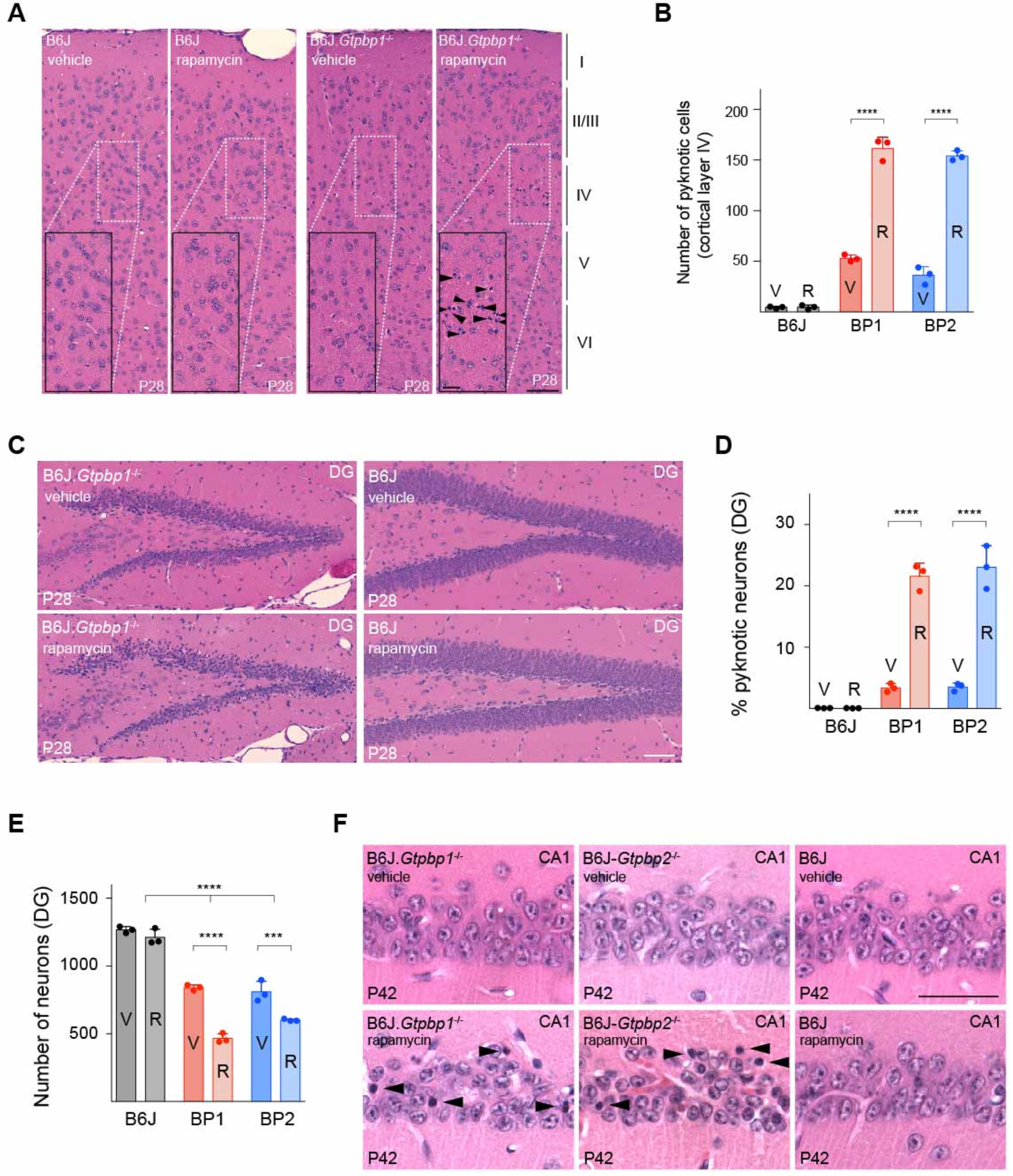
Inhibition of mTOR signaling accelerates cell death in B6J.*Gtpbp1*^-/-^ and B6J-*Gtpbp2*^-/-^ mice. (*A*) Sagittal sections of the cortex of P28 mice injected with vehicle or rapamycin for two weeks stained with hematoxylin and eosin. Higher magnification images of layer IV (rectangle) are shown for each genotype. Cortical layers are indicated by Roman numerals and arrowheads indicate pyknotic cells. (*B*) Number of pyknotic cells in layer IV of the cortex in (B6J), BP1 (B6J.*Gtpbp1*^-/-^), and BP2 (B6J-*Gtpbp2*^-/-^) mice injected with vehicle (V) or rapamycin (R). Data represent mean ± SD. (*C*) Hematoxylin and eosin stained sagittal sections of the hippocampus of P28 mice injected with vehicle or rapamycin for 14 days. Arrowheads indicate pyknotic cells. Note: Due to the large number of pyknotic cells in rapamycin-treated B6J.*Gtpbp1*^-/-^ mice, arrowheads to indicate pyknotic cells were not included. (*D*) Percent of pyknotic neurons of the dentate gyrus (DG) in vehicle (V) or rapamycin (R) treated control (B6J), BP1 (B6J.*Gtpbp1*^-/-^), and BP2 (B6J-*Gtpbp2*^-/-^) mice. Data represent mean ± SD. (*E*) Total number of neurons of the dentate gyrus (DG) in vehicle (V) or rapamycin (R) treated control (B6J), BP1 (B6J.*Gtpbp1*^-/-^), and BP2 (B6J-*Gtpbp2*^-/-^) mice. Data represent mean ± SD. (*F*) Sagittal sections of the hippocampus of P42 mice injected with vehicle or rapamycin for two weeks stained with hematoxylin and eosin. Arrowheads indicate pyknotic cells. Scale bars: 50μm and 20μm (higher magnification) (*A*); 50μm (*C, F*). Two-way ANOVA, corrected for multiple comparisons using Tukey method (*B, D, E*). *** *P* ≤ 0.001, **** *P* ≤ 0.0001.

## Discussion

Ribosome speed during translation is regulated to allow proper folding of the nascent peptide and functional protein production. However, prolonged pausing, or stalling of translating ribosomes can be detrimental to cells. Thus cells have evolved multiple quality control mechanisms that mediate steps from recognition of stalled ribosomes to the resolution of these complexes (Brandman and Hegde, 2016; Joazeiro, 2019). We previously identified GTPBP2 as ribosome rescue factor essential for neuronal survival during tRNA deficiency (Ishimura et al., 2014). The high homology of GTPBP1 and GTPBP2, their broad expression patterns, and the lack of overt phenotypes of *Gtpbp1*-deficient mice on a mixed genetic background, had suggested functional redundancy of these translational GTPases (Senju et al., 2000). However, we show here that loss of *Gtpbp1* leads to widespread neurodegeneration in B6J mice. As observed in *Gtpbp2*^-/-^ mice, neuron loss in *Gtpbp1*^-/-^ mice is dependent on a mutation in B6J strain that disrupts *n-Tr20* tRNA^Arg^_UCU_ processing (Ishimura et al., 2014). Ribosome profiling of the *Gtpbp1*^-/-^ cerebellum revealed increased ribosome occupancy at AGA codons as we previously observed for *Gtpbp2*^-/-^ mice (Ishimura et al., 2014). These data suggest that both GTPBP1 and GTPBP2 function as ribosome rescue factors and are essential genes in many neurons when *n-Tr20* levels are diminished. In addition, the lack of neurodegeneration in mice heterozygous for mutations in both *Gtpbp1* and *Gtpbp2* and the lack of an additive phenotype in the brain of mice lacking both genes, suggest that these trGTPases function in the same pathways to mitigate ribosome stalling, perhaps mediating different steps in this pathway.

GTPBP1 and GTPBP2 share domain homology with other trGTPases including the yeast protein Hbs1 (Hbs1L in mammals) (Atkinson, 2015). Hbs1 and its interacting partner Dom34 recognize paused ribosomes on the 3’end of truncated mRNAs and in the 3’UTR of mRNAs, and resolution of paused ribosomes is in part gated by the GTPase activity of Hbs1 (Becker et al., 2012; Guydosh et al., 2017; Guydosh and Green, 2014; Juszkiewicz et al., 2020; Pisareva et al., 2011; Shoemaker and Green, 2011). Recent biochemical studies revealed that the GTPase activity of GTPBP1 and GTPBP2 is not stimulated in the presence of 80S ribosomes (Zinoviev et al., 2018), suggesting that these two trGTPases likely function differently from that of Hbs1. In agreement, Dom34/Hbs1 did not mediate pause resolution in a codon-specific manner when pausing was induced at His codons upon inhibition of histidine biosynthesis (Guydosh and Green, 2014).

During the process of pause resolution, multiple factors are recruited to mediate ribosome recycling, degradation of the nascent peptide chains, or mRNAs associated with stalled ribosomes (Collart and Weiss, 2019; D’Orazio et al., 2019; Ibrahim et al., 2018; Pelechano et al., 2015). Interestingly, GTPBP1 can stimulate exosomal degradation of mRNAs (Woo et al., 2011; Zinoviev et al., 2018) and thus may provide a link between elongation defects and mRNA degradation. Although we did not observe a strong correlation between mRNA abundance and ribosome pausing in the *Gtpbp1*^-/-^ cerebellum, the stochastic nature of ribosome stalling could make it difficult to observe the corresponding, similarly stochastic changes in mRNA decay. RNA sequencing techniques that are specifically designed to capture degradation intermediates may be more efficient to investigate mRNA degradation *in vivo* (Ibrahim et al., 2018).

As we previously observed in the B6J-*Gtpbp2*^-/-^ cerebellum (Ishimura et al., 2016), loss of *Gtpbp1* activated the ISR via activation of GCN2, and deletion of GCN2 in B6J.*Gtpbp1*^-/-^ mice resulted in accelerated, and more widespread, neurodegeneration. GCN2 can be activated by uncharged tRNA that accumulates during amino acid deprivation (Masson, 2019). However, increases in uncharged tRNA were not observed in the B6J-*Gtpbp2*^-/-^ cerebellum, suggesting other mechanisms underlie GCN2 activation (Ishimura et al., 2016). Indeed, recent biochemical and genetic studies revealed that the P-stalk, a pentameric complex located near the A-site of the ribosome, interacts with GCN2 and activates it, suggesting that translation elongation defects may be sensed directly by GCN2 (Harding et al., 2019; Inglis et al., 2019). Activation of the ISR and its accompanying protective function was observed across multiple neuronal populations in *Gtpbp1*^-/-^ and *Gtpbp2*^-/-^ mice. Interestingly, expression of some ATF4 target genes varied between types of neurons, suggesting that additional cell-type specific mechanisms, such as those controlling formation of ATF4 heterodimers, post-translational modifications of ATF4, or epigenetic regulation of target genes may modulate expression of these genes (Wortel et al., 2017).

We also show that defects in ribosome pause resolution in the B6J.*Gtpbp1*^-/-^ and B6J-*Gtpbp2*^-/-^ brain are associated with decreased mTORC1 signaling. In contrast to ISR activation, which prolonged neuronal survival, inhibition of mTOR signaling negatively affected neuronal survival in mutant mice, despite the fact that the observed changes in both of these pathways are associated with decreased translation initiation. Previous studies showed that during amino acid starvation of HEK293T cells, reduction of mTOR activity and translation correlated with a reduction in ribosome pausing (Darnell et al., 2018). While inhibition of mTORC1 may be beneficial at the level of translation in some instances of ribosome stress, the negative impact we observe on neuronal survival may reside in the severity or the duration of the stressor and/or the influence of mTORC1 on other cellular processes including ribosomal RNA processing, gene transcription, protein degradation, ribosomal and mitochondrial biogenesis that could inflict additional obstructions (Laplante and Sabatini, 2013, 2009; López et al., 2019; Mayer and Grummt, 2006; Puertollano, 2014).

The dramatic downregulation of mTORC1 activity in granule cells of the B6J.*Gtpbp1*^-/-^ and B6J-*Gtpbp2*^-/-^ dentate gyrus relative to other neurons suggests that the cellular context plays a role in modulating this signaling pathway upon ribosomal stress. A previous study observed increased mTOR activity in epidermal stem cells upon loss of the ribosome recycling factor *Pelo*, and suppression of mTOR signaling partially restored cellular defects *in vivo* (Liakath-Ali et al., 2018). We recently observed that the relatively low levels of ribosome pausing that occur upon tRNA deficiency (with normal levels of GTPBP1 and GTPBP2) can lead to mTORC1 inhibition and alter neuronal physiology (Kapur et al., 2020). Together, these data demonstrate that elongation defects may lead to hyper- or hypoactivity of mTOR signaling and these changes in mTORC1 appear to negatively modulate cellular homeostasis and survival.

How mTORC1 responds to translational stress, particularly the mechanism underlying its cell type-specific response, is unknown. Proteomic experiments have revealed mTOR-dependent phosphorylation of translation initiation factors and ribosomal proteins. Interestingly, multiple phosphorylation sites were observed on the surface of the 80S ribosome, suggesting that mTOR or the mTOR-associated kinases directly access the ribosome (Jiang et al., 2016). The interaction of mTOR and/or its downstream kinases with the ribosome may provide a mechanism for mTOR to simultaneously monitor and regulate translation. Because protein synthesis varies between cell types and changes during development (Blair et al., 2017; Buszczak et al., 2014; Castelo-Szekely et al., 2017; Gonzalez et al., 2014; Sudmant et al., 2018), the rate of translation may differentially impact elongation defects and thus the signaling pathways which control translation.

## Material and Methods

### Mice strains

*Gtpbp1*^-/+^ mice were generated previously on a mixed genetic background with a portion of exon 3 and all of exon 4 replaced with a PGK-Neo cassette (Senju et al., 2000). These mice were backcrossed to C57BL/6J mice for more than 10 generations to generate congenic B6J.*Gtpbp1*^-/-^ mice and genotyped with the following primers (wild type forward primer: 5’GAGTACGGGCTGAGTGAAGC3’, wild type reverse primer: 5’TGGACAGGAACCTGATGTGA3’, mutant forward primer: 5’TACGCCACCGTGAAGAGCAT3’, mutant reverse primer: 5’AGGGGAGGAGTGGAAGGTGG3’). Homozygosity for the tRNA (*n-Tr20*^J/J^) mutation was confirmed by genotyping (Ishimura et al., 2014). For transgene rescue experiments, B6J.Tg(*n-Tr20*^N/N^); *Gtpbp1*^-/-^ mice were generated by crossing B6J.*Gtpbp1*^+/-^ mice to B6J-Tg(*n-Tr20*^N/N^) mice that transgenically express wild type levels of wild type *n-Tr20* (Ishimura et al., 2014) and then backcrossing to B6J.*Gtpbp1*^-/+^ mice. The generation of B6J-Tg(*n-Tr20*^N/N^) and B6J-*nmf205*^-/-^ (B6J.*Gtpbp2*^-/-^) has been previously described (Ishimura et al., 2014). C57BL/6J (B6J) and B6J.*Gcn2*^-/-^ mice (*Eif2ak4^tm1.2Dron^*) were obtained from The Jackson Laboratory. Mice of both sexes were used for experiments. The Jackson Laboratory Animal Care and Use Committee and The University of California San Diego Animal Care and Use Committee approved all mouse protocols.

### Rapamycin treatment

The rapamycin (LC laboratories) stock solution (50mg/ml) was prepared in ethanol and diluted on the day of injection in equal volumes of a 10 % PEG-400/8% ethanol solution and 10% Tween-80. Mice were injected intraperitoneally with 5mg/kg rapamycin or vehicle daily; injections were performed from P14-P21 for tissue collection (RNA isolation or western blotting) or from P14-P28 or P28-P42 for histological analysis.

### Histology and immunohistochemistry

Anesthetized mice were transcardially perfused with 4% paraformaldehyde (PFA) for immunofluorescence, 10% neutral buffered formalin (NBF), for *in situ* hybridization or immunofluorescence, or Bouin’s fixative for histology. Tissues were post-fixed overnight and embedded in paraffin. For histological analysis, sections were deparaffinized, rehydrated, and were stained with hematoxylin and eosin according to standard procedures. Histological slides were imaged using a digital slide scanner (Hamamatsu).

For quantification of cerebellar granule cells, the total number of granule cells (viable and pyknotic cells) were counted in a 0.02mm^2^ area from lobule VI or IX and averaged from three sections per brain spaced 100μm apart at midline, and averaged. For quantification of hippocampal neurons, the number of neurons with, and without, pyknotic nuclei were counted in the CA1 or dentate gyrus of the hippocampus and averaged from three sections, spaced 50-70μm apart and about 1.5mm off midline, per brain. For quantification of pyknotic nuclei in the cortex, the total number of pyknotic nuclei were counted across the entire section and averaged from three sections per animal spaced 50-70μm apart and about 2-2.5mm off midline, per brain. All histological quantifications were performed with three mice of each genotype and time point using mice of both sexes.

For immunofluorescence, antigen retrieval on deparaffinized PFA-fixed sections was performed by microwaving sections in 0.01M sodium citrate buffer (pH 6.0, 0.05% Tween-20), three times for 3 minutes each. NBF-fixed sections were microwaved for three times for 3 minutes, followed by two times for 9 minutes (NBF-fixed sections). PFA- or NBF-fixed sections were incubated with the following primary antibodies: rabbit anti-cleaved caspase 3-D175 (Cell Signaling, #9661, 1:100, PFA-fixed tissue), Ctip2 (Abcam, ab18465, 1:500, PFA-fixed tissue), p-S6^240/244^ (Cell Signaling, #5364, 1:1000, NBF-fixed tissue), and RPS6 (Santa Cruz Biotechnology, sc-74459, 1:500, NBF-fixed tissue). Detection of primary antibodies was performed with goat anti-mouse Alexa Fluor-488, goat anti-rabbit Alexa Fluor-488, goat anti-rat Alexa Fluor-555, and goat antirabbit Alexa Fluor-555 secondary antibodies (Invitrogen). Sections were counterstained with DAPI, and treated with Sudan black to quench autofluorescence.

### RNAscope (*in situ* hybridization)

*In situ* hybridization of *Gtpbp1, Gtpbp2, Sesn2, Slc7a1, Ddr2*, and *Chac1* probes was performed as per the manufacturer’s protocol (RNAscope Multiplex Fluorescent Reagent Kit v2; Advanced Cell Diagnostics). Briefly, deparaffinized NBF-fixed sections were treated for 15 minutes with Target Retrieval Reagent at 100°C and subsequently, treated with Protease Plus for 30 minutes at 40°C. RNAscope probes were hybridized for 2 hours. TSA^®^ Plus Cyanine 5 (PerkinElmer, 1:1,000) was used as a secondary fluorophore for C1 probes (*Gtpbp2, Slc7a1, Chac1*) and TSA^®^ Plus Cyanine 3 (PerkinElmer, 1:2,000) was used as a secondary fluorophore for C2 probes (*Gtpbp1, Sesn2, Ddr2*).

### Reverse transcription, quantitative PCR, and genomic PCR analysis

Whole brain, cerebella or hippocampi were isolated and immediately frozen in liquid nitrogen. Total RNA was extracted with Trizol reagent (Life Technologies). cDNA synthesis was performed on DNase-treated (DNA-free DNA Removal Kit, Life Technologies AM1906) total RNA using oligo(dT) primers and the SuperScript III First-Strand Synthesis System (Life Technologies). Quantitative RT-PCR (qRT) reactions were performed using iQ SYBR Green Supermix (Bio-Rad) and an CFX96 Real-Time PCR Detection System (Bio-Rad). Reactions were performed with primers previously published (Ishimura et al., 2016). Expression levels of the genes of interest were normalized to β-actin (forward primer: 5’GGCTGTATTCCCCTCCATCG3’, reverse primer: 5’ CCAGTTGGTAACAATGCCATGT3’) using the 2^-ΔΔCT^ method (Livak and Schmittgen, 2001) and expressed as fold change ± standard error of the mean (SEM) relative to control (B6J).

### RNA-Seq library preparation

Cerebella from various strains were isolated from 3-week-old mice (P21, n=3 mice for each genotype) and immediately frozen in liquid nitrogen. RNA-Seq libraries were prepared using the TruSeq v2 mRNA kit (Illumina). Briefly, total cerebellar RNA was isolated and RNA quality was assessed on Agilent TapeStation. mRNA was purified using biotin-tagged polydT oligonucleotides and streptavidin-coated magnetic beads. After fragmentation of mRNA, libraries were prepared according to manufacturer’s instructions. Paired end reads (2×100bp) were obtained using the HiSeq 4000 (Illumina).

### Ribosome profiling library construction

Ribosome profiling libraries were generated as previously described (Ingolia et al., 2012; Ishimura et al., 2014) with some minor modifications. Briefly, dissected cerebella were immediately frozen in liquid nitrogen. One cerebellum from P21 mice was used for each biological replicate, and three biological replicates were prepared for each genotype. Tissue homogenization was performed with a mixer mill (Retsch MM400) in 350μl lysis buffer (20mM Tris-Cl, pH 8.0, 150 mM NaCl, 5mM MgCl_2_, 1mM DTT, 100μg/ml CHX, 1% (v/v) TritonX-100, 25units/ml Turbo DNaseI). RNase I-treated lysates were overlaid on top of a sucrose cushion in 5ml Beckman Ultraclear tubes and centrifuged in an SW55Ti rotor for 4 hours at 4°C at 46,700 rpm. Pellets were resuspended and RNA was extracted using the miRNeasy kit (Qiagen) according to manufacturer’s instructions. 26-34 nucleotide RNA fragments were purified by electrophoresis on a denaturing 15% gel. Linker addition, cDNA generation (first-strand synthesis was performed at 50°C for 1h), circularization, rRNA depletion, and amplification of cDNAs with indexing primers were performed. Library quality and concentration was assessed using high sensitivity D1000 screen tape on the Agilent TapeStation, Qubit 2.0 Fluorometer, and qPCR. Libraries were run on HiSeq4000 (SR75).

### RNA-Seq data analysis

Reads were quantified using kallisto version 0.42.4 (Bray et al., 2016) and pseudo-aligned to Gencode M20 transcripts reference. Differential expression was performed using sleuth version 0.30.0 (Pimentel et al., 2017).

### Ribosome profiling data analysis

Reads were clipped to remove adaptor sequences (CTGTAGGCACCATCAAT) using fastx_clipper and trimmed so that reads start on the second base using fastx_trimmer (http://hannonlab.cshl.edu/fastx_toolkit/). Reads containing ribosomal RNA were then filtered out by mapping to a ribosomal RNA reference using bowtie2 version 2.2.3 using parameters -L 13 (Langmead and Salzberg, 2012). Remaining reads were mapped to a mm10 mouse reference using a Gencode M20 annotation, or a Gencode M20 transcript reference using hisat2 version 2.1.0 (Kim et al., 2019). Ribosomal A-sites were identified using RiboWaltz version 1.0.1 (Lauria et al., 2018), and read lengths 29-33 were retained for further analysis. Observed/expected reads were calculated for each transcript with alignments with the expected reads being the read density expected at a given site with a given codon, assuming that reads are uniformly distributed across the coding part of the transcript. Pauses were identified using previous methodology (Ishimura et al., 2014) using a 0.5 reads/codon in all samples to threshold transcripts to analyze. Correlation between gene pauses and gene expression was calculated by taking the average pause scores of transcripts associated with a gene for transcripts with ≥ 0.5 reads/codon across the B6J.*Gtpbp1*^-/-^ samples. The sum of these pause scores across all replicates was then correlated with the sleuth *beta* values (in the B6J.*Gtpbp1*^-/-^ vs. B6J comparison) for genes that passed the reads/codon thresholding in the ribosome profiling datasets.

Read counts for translational efficiency (TE) analysis were quantified using featureCounts (Liao et al., 2014) with footprints overlapping CDS features and RNA read pairs overlapping gene exon features. Histone mRNAs were removed from the analysis and TE was quantified and tested for using riborex version 2.3.4 using the DESeq2 engine (Li et al., 2017). TE analysis of AGA A-site filtered datasets was performed by identifying AGA codons in the transcriptome and transferring the coordinates to the genome using ensembldb v.2.6.8 (Rainer et al., 2019). Then reads with AGAs in the A-site (±1 codon based on riboWaltz A-site offset for each read length) were removed and the above TE analysis was performed. Sets of mouse transcripts with known TOP motifs were identified in (Yamashita et al., 2008).

### Gene ontology (GO) and pathway analysis

RNA-sequencing data were analyzed using Ingenuity Pathway Analysis (QIAGEN Inc., https://www.qiagenbioinformatics.com/products/ingenuity-pathway-analysis). Gene Ontology (GO) pathway analysis was performed using the DAVID bioinformatics web server (http://david.abcc.ncifcrf.gov) by uploading the gene lists from our ribosome profiling analysis (AGA stalling transcripts, z ≥ 10, stalls detected in any biological replicates). The functional annotation chart and clustering analysis modules were utilized for gene-term enrichment analysis, and terms with a Benjamini-Hochberg adjusted p-value ≤ 0.05 were considered enriched.

### Western blotting

Cerebella or hippocampi were isolated and immediately frozen in liquid nitrogen. Proteins were extracted by homogenizing tissue in 5 volumes of RIPA buffer with cOmplete Mini, EDTA-free Protease inhibitor Cocktail (Roche), sonicated two times for 10 seconds (Branson, 35% amplitude), incubated for 30 minutes at 4°C, centrifuged at 16’000xg for 25 minutes, and 25μg of whole protein lysate were resolved on SDS-PAGE gels prior to transfer to PVDF membranes (GE Healthcare Life Sciences, #10600023) using a tank blotting apparatus (BioRad).

For detection of phosphoproteins, frozen tissue samples were homogenized in 5 volumes of homogenization buffer (50mM Hepes/KOH, pH 7.5, 140mM potassium acetate, 4mM magnesium acetate, 2.5mM dithiothreitol, 0.32M sucrose, 1mM EDTA, 2mM EGTA) (Carnevalli et al., 2004), supplemented with phosphatase and protease inhibitors (PhosStop and cOmplete Mini, EDTA-free Protease inhibitor Cocktail, Roche). Lysate samples were centrifuged at 12,000xg for 7 minutes, and either 30μg (detection of ribosomal protein S6) or 70μg (detection of eIF2αlpha) of whole protein lysate were resolved on SDS-PAGE gels prior to transfer to PVDF membranes. After blocking in 5% nonfat dry milk (Cell Signaling, #9999S), blots were probed with primary antibodies at 4°C overnight: rabbit anti-phsopho-S6^240/244^ (Cell Signaling, #5364, 1:4000), mouse anti-RPS6 (Santa Cruz Biotechnology, sc-74459, 1:2000), rabbit anti-phospho-eIF2αlpha^S51^ (Cell Signaling, #9721, 1:1000), rabbit anti-EIF2alpha (Cell Signaling, #9722, 1:2000), mouse anti-Vinculin (Sigma, V-9131, 1:20,000), and followed by incubation with HRP-conjugated secondary antibodies for 2 hours at room temperature: goat anti-rabbit IgG (BioRad, #170-6515) or goat antimouse IgG (BioRad, #170-6516). Signals were detected with SuperSignal West Pico Chemiluminescent substrate (ThermoScientific, #34080).

### Statistics

For quantification of protein expression, RNA expression, or histological quantifications, p-values were computed in GraphPad Prism using either multiple *t*-tests, one-way ANOVA, or two-way ANOVA, and corrected for multiple comparisons as indicated in the figure legends. All quantifications were performed with at least three mice of each genotype and time point using mice of either sex.

## Acknowledgements

We thank T. Jucius, and A. Kano for technical assistance, Drs. Satori Senju and Yasuharu Mishimura for providing *Gtpbp1*^-/-^ mice, the IGM Genomics Center at the University of California San Diego (UCSD) for support with the preparation of RNA sequencing libraries and sequencing, and the UCSD School of Medicine Microcopy Core for providing access to microscopy equipment (Grant P30 NS047101). This work was supported by NIH NS094637 (SLA). SLA is an investigator of the Howard Hughes Medical Institute.

## Author Contributions

MT and SLA designed experiments and wrote the paper. ALG performed RNAscope and immunofluorescence experiments for ribosomal protein S6. TD and MT performed experiments using GCN2 mice. RI performed initial phenotype analysis of *Gtpbp1*^-/-^ mice. MT performed mouse and molecular biology experiments under SLA’s guidance. SIA performed the computational analysis of RNA-sequencing and ribosome profiling data under JHC’s guidance.

## References

Atkinson GC. 2015. The evolutionary and functional diversity of classical and lesser-known cytoplasmic and organellar translational GTPases across the tree of life. BMC Genomics 16:78. doi:10.1186/s12864-015-1289-7

Becker T, Franckenberg S, Wickles S, Christopher J, Anger AM, Armache J, Sieber H, Berninghausen O, Daberkow I, Karcher A, Thomm M, Hopfner K, Green R, Beckmann R. 2012. Structural Basis of Highly Conserved Ribosome Recycling in Eukaryotes and Archaea. Nature 482:501–506. doi:10.1038/nature10829

Bertoli-Avella AM, Garcia-Aznar JM, Brandau O, Al-Hakami F, Yüksel Z, Marais A, Grüning NM, Abbasi Moheb L, Paknia O, Alshaikh N, Alameer S, Marafi MJ, Al-Mulla F, Al-Sannaa N, Rolfs A, Bauer P. 2018. Biallelic inactivating variants in the GTPBP2 gene cause a neurodevelopmental disorder with severe intellectual disability. Eur J Hum Genet 26:592–598. doi:10.1038/s41431-018-0097-3

Blair JD, Hockemeyer D, Doudna JA, Bateup HS, Floor SN. 2017. Widespread translational remodeling during human neuronal differentiation. Cell Rep 21:2005–2016. doi:10.1016/j.celrep.2017.10.095

Brandman O, Hegde RS. 2016. Ribosome-associated Protein Quality Control. Nat Struct Mol Biol 23:7–15. doi:10.1038/nsmb.3147

Bray NL, Pimentel H, Melsted P, Lior Pachter. 2016. Near-optimal Probabilistic RNA-seq Quantification. Nat Biotechnol 34:525–527. doi:10.1038/nbt.3519

Brule CE, Grayhack EJ. 2017. Synonymous Codons: Choose Wisely for Expression. Trends Genet 33:283–297. doi:10.1016/j.tig.2017.02.001

Buhr F, Jha S, Thommen M, Mittelstaet J, Kutz F, Schwalbe H, Rodnina M V, Komar AA. 2016. Synonymous Codons Direct Cotranslational Folding Toward Different Protein Conformations. Mol Cell 61:341–351. doi:10.1016/j.molcel.2016.01.008

Buszczak M, Signer RAJ, Morrison SJ. 2014. Cellular differences in protein synthesis regulate tissue homeostasis. Cell 159:242–251. doi:doi:10.1016/j.cell.2014.09.016.

Carnevalli LS, Pereira CM, Longo BM, Jaqueta CB, Avedissian M, Mello LEAM, Castilho BA. 2004. Phosphorylation of translation initiation factor eIF2α in the brain during pilocarpine-induced status epilepticus in mice. Neurosci Lett 357:191–194. doi:10.1016/j.neulet.2003.12.093

Carter MT, Venkateswaran S, Shapira-Zaltsberg G, Davila J, Humphreys P, Kernohan KD, Boycott KM. 2019. Clinical delineation of GTPBP2-associated neuro-ectodermal syndrome: Report of two new families and review of the literature. Clin Genet 95:601–606. doi:10.1111/cge.13523

Castelo-Szekely V, Arpat AB, Janich P, Gatfield D. 2017. Translational contributions to tissue specificity in rhythmic and constitutive gene expression. Genome Biol 18:1–17. doi:10.1186/s13059-017-1222-2

Chaney JL, Clark PL. 2015. Roles for Synonymous Codon Usage in Protein Biogenesis. Annu Rev Biophys 44:143–166. doi:10.1146/annurev-biophys-060414-034333

Chesnokova E, Bal N, Kolosov P. 2017. Kinases of eIF2α switch translation of mRNA subset during neuronal plasticity. Int J Mol Sci 18:2213. doi:10.3390/ijms18102213

Chu D, Kazana E, Bellanger N, Singh T, Tuite. MF, Haar T von der. 2014. Translation elongation can control translation initiation on eukaryotic mRNAs. EMBO J 33:21–34. doi:10.1002/embj.201385651

Collart MA, Weiss B. 2019. Ribosome pausing, a dangerous necessity for co-translational events. Nucleic Acids Res 48:1043–1055. doi:10.1093/nar/gkz763

D’Orazio KN, Wu CCC, Sinha N, Loll-Krippleber R, Brown GW, Green R. 2019. The endonuclease cue2 cleaves mRNAs at stalled ribosomes during no go decay. Elife 8:1–27. doi:10.7554/eLife.49117.001

Dai M, Lu H. 2009. Cross talk between c-myc and ribosomes. J Cell Biochem 105:670–677. doi:10.1002/jcb.21895.Crosstalk

Dalton LE, Healey E, Irving J, Marciniak SJ. 2012. Phosphoproteins in Stress-Induced Disease. Prog Mol Biol Transl Sci 106:189–221. doi:10.1016/B978-0-12-396456-4.00003-1

Darnell A, Subramaniam A, O’Shea E. 2018. Translational control through differential ribosome pausing during amino acid limitation in mammalian cells. Mol Cell 71:229–243. doi:10.1016/j.molcel.2018.06.041

Drummond DA, Wilke CO. 2008. Mistranslation-Induced Protein Misfolding as a Dominant Constraint on Coding-Sequence Evolution. Cell 134:341–352. doi:10.1016/j.cell.2008.05.042

Gonzalez C, Sims JS, Hornstein N, Mela A, Garcia F, Lei L, Gass DA, Amendolara B, Bruce JN, Canoll P, Sims PA. 2014. Ribosome profiling reveals a cell-type-specific translational landscape in brain tumors. J Neurosci 34:10924–10936. doi:10.1523/JNEUROSCI.0084-14.2014

Guydosh NR, Green R. 2014. Dom34 rescues ribosomes in 3’ untranslated regions. Cell 156:950–962. doi:10.1016/j.cell.2014.02.006

Guydosh NR, Kimmig P, Walter P, Green R. 2017. Regulated Ire1-dependent mRNA decay requires no-go mRNA degradation to maintain endoplasmic reticulum homeostasis in S. Pombe. Elife 6:e29216. doi:10.7554/eLife.29216

Harding HP, Ordonez A, Allen F, Parts L, Inglis AJ, Williams RL, Ron D. 2019. The ribosomal P-stalk couples amino acid starvation to GCN2 2 activation in mammalian cells. Elife 8:1–19. doi:10.7554/eLife.50149

Harding HP, Zhang Y, Zeng H, Novoa I, Lu PD, Calfon M, Sadri N, Yun C, Popko B, Paules R, Stojdl DF, Bell JC, Hettmann T, Leiden JM, Ron D. 2003. An integrated stress response regulates amino acid metabolism and resistance to oxidative stress. Mol Cell 11:619–633. doi:10.1016/S1097-2765(03)00105-9

Ibrahim F, Maragkakis M, Alexiou P, Mourelatos Z. 2018. Ribothrypsis, a novel process of canonical mRNA decay, mediates ribosome-phased mRNA endonucleolysis. Nat Struct Mol Biol 25:302–310. doi:10.1038/s41594-018-0042-8

Inglis AJ, Masson GR, Shao S, Perisic O, McLaughlin SH, Hegde RS, Williams RL. 2019. Activation of GCN2 by the ribosomal P-stalk. Proc Natl Acad Sci U S A 116:4946–4954. doi:10.1073/pnas.1813352116

Ingolia NT, Brar GA, Rouskin S, McGeachy AM, Weissman JS. 2012. The ribosome profiling strategy for monitoring translation in vivo by deep sequencing of ribosome-protected mRNA fragments. Nat Protoc 7:1534–1550. doi:10.1038/nprot.2012.086

Ishimura R, Nagy G, Dotu I, Chuang JH, Ackerman SL. 2016. Activation of GCN2 kinase by ribosome stalling links translation elongation with translation initiation. Elife 5:1–22. doi:10.7554/eLife.14295

Ishimura R, Nagy G, Dotu I, Zhou H, Yang XL, Schimmel P, Senju S, Nishimura Y, Chuang JH, Ackerman SL. 2014. Ribosome stalling induced by mutation of a CNS-specific tRNA causes neurodegeneration. Science (80-) 345:455–459. doi:10.1126/science.1249749

Jaberi E, Rohani M, Shahidi GA, Nafissi S, Arefian E, Soleimani M, Rasooli P, Ahmadieh H, Daftarian N, KaramiNejadRanjbar M, Klotzle B, Fan J-B, Turk C, Steemers F, Elahi E. 2016. Identification of Mutation in GTPBP2 in Patients of a Family With Neurodegeneration Accompanied by Iron Deposition in the Brain. Neurobiol Aging 38:216.e11–216.e18. doi:10.1016/j.neurobiolaging.2015.10.034

Jaeseok Han, Back SH, Hur J, Lin Y-H, Gildersleeve R, Shan J, Yuan CL, Krokowski D, Wang S, Hatzoglou M, Kilberg MS, Sartor MA, Kaufman RJ. 2013. ER-stress-induced transcriptional regulation increases protein synthesis leading to cell death. Nat Cell Biol 15:481–490. doi:10.1038/ncb2738

Jiang X, Feng S, Chen Y, Feng Y, Deng H. 2016. Proteomic analysis of mTOR inhibition-mediated phosphorylation changes in ribosomal proteins and eukaryotic translation initiation factors. Protein Cell 7:533–537. doi:10.1007/s13238-016-0279-0

Joazeiro CAP. 2019. Mechanisms and Functions of Ribosome-Associated Protein Quality Control. Nat Rev Mol Cell Biol 20:368–383. doi:10.1038/s41580-019-0118-2

Juszkiewicz S, Speldewinde SH, Wan L, Svejstrup JQ, Hegde RS. 2020. The ASC-1 Complex Disassembles Collided Ribosomes. Mol Cell 79:603–614. doi:10.1016/j.molcel.2020.06.006

Kaiser CM, Liu K. 2018. Folding Up and Moving on-Nascent Protein Folding on the Ribosome. J Mol Biol 430:4580–4591. doi:10.1016/j.jmb.2018.06.050

Kapur M, Ganguly A, Nagy G, Adamson SI, Chuang JH, Frankel WN, Ackerman SL. 2020. Expression of the Neuronal tRNA n-Tr20 Regulates Synaptic Transmission and Seizure Susceptibility Article Expression of the Neuronal tRNA n-Tr20 Regulates Synaptic Transmission and Seizure Susceptibility. Neuron. doi:10.1016/j.neuron.2020.07.023

Kim D, Paggi JM, Park C, Bennett C, Salzberg SL. 2019. Graph-based genome alignment and genotyping with HISAT2 and HISAT-genotype. Nat Biotechnol 37:907–915. doi:10.1038/s41587-019-0201-4

Langmead B, Salzberg SL. 2012. Fast gapped-read alignment with Bowtie 2. Nat Methods 9:357–359. doi:10.1038/nmeth.1923

Laplante M, Sabatini DM. 2013. Regulation of mTORC1 and its impact on gene expression at a glance. J Cell Sci 126:1713–1719. doi:10.1242/jcs.125773

Laplante M, Sabatini DM. 2009. mTOR signaling at a glance. J Cell Sci 122:3589–3594. doi:10.1242/jcs.051011

Lauria F, Tebaldi T, Bernabò P, Groen EJN, Gillingwater TH, Viero G. 2018. riboWaltz: Optimization of ribosome P-site positioning in ribosome profiling data. PLoS Comput Biol 14:1–20. doi:10.1371/journal.pcbi.1006169

Li W, Wang W, Uren PJ, Penalva LOF, Smith AD. 2017. Riborex: Fast and flexible identification of differential translation from Ribo-seq data. Bioinformatics 33:1735–1737. doi:10.1093/bioinformatics/btx047

Liakath-Ali K, Mills EW, Sequeira I, Lichtenberger BM, Pisco AO, Sipilä KH, Mishra A, Yoshikawa H, Wu CC-C, Tony Ly AIL, Adham IM, Green R, Watt FM. 2018. An Evolutionarily Conserved Ribosome-Rescue Pathway Maintains Epidermal Homeostasis. Nature 556:376–380. doi:10.1038/s41586-018-0032-3

Liao Y, Smyth GK, Shi W. 2014. FeatureCounts: An efficient general purpose program for assigning sequence reads to genomic features. Bioinformatics 30:923–930. doi:10.1093/bioinformatics/btt656

Livak KJ, Schmittgen TD. 2001. Analysis of relative gene expression data using real-time quantitative PCR and the 2-ΔΔCT method. Methods 25:402–408. doi:10.1006/meth.2001.1262

López KG de la C, Mariel Esperanza Toledo Guzmán EOS, Carrancá AG. 2019. mTORC1 as a Regulator of Mitochondrial Functions and a Therapeutic Target in Cancer. Front Oncol 9:1–22. doi:10.3389/fonc.2019.01373

Ma XM, Blenis J. 2009a. mTOR complex 2 signaling and functions. Nat Rev Mol Cell Biol 10:2305–2316. doi:10.4161/cc.10.14.16586

Ma XM, Blenis J. 2009b. Molecular mechanisms of mTOR-mediated translational control. Nat Rev Mol Cell Biol 10:307–318. doi:10.1038/nrm2672

Masson GR. 2019. Towards a model of GCN2 activation. Biochem Soc Trans 47:1481–1488. doi:10.1042/BST20190331

Mayer C, Grummt I. 2006. Ribosome biogenesis and cell growth: mTOR coordinates transcription by all three classes of nuclear RNA polymerases. Oncogene 25:6384–6391. doi:10.1038/sj.onc.1209883

Nandagopal N, Roux PP. 2015. Regulation of global and specific mRNA translation by the mTOR signaling pathway. Translation 3:e983402. doi:10.4161/21690731.2014.983402

Park Y, Reyna-Neyra A, Philippe L, Thoreen CC. 2017. mTORC1 Balances Cellular Amino Acid Supply with Demand for Protein Synthesis through Post-transcriptional Control of ATF4. Cell Rep 19:1083–1090. doi:10.1016/j.celrep.2017.04.042

Pelechano V, Wei W, Steinmetz LM. 2015. Widespread co-translational RNA decay reveals ribosome dynamics. Cell 161:1400–1412. doi:10.1016/j.cell.2015.05.008

Pimentel H, Bray NL, Puente S, Melsted P, Pachter L. 2017. Differential Analysis of RNA-seq Incorporating Quantification Uncertainty. Nat Methods 14:687–690. doi:10.1038/nmeth.4324

Pisareva VP, Skabkin MA, Hellen CUT, Pestova T V., Pisarev A V. 2011. Dissociation by Pelota, Hbs1 and ABCE1 of mammalian vacant 80S ribosomes and stalled elongation complexes. EMBO J 30:1804–1817. doi:10.1038/emboj.2011.93

Puertollano R. 2014. mTOR and lysosome regulation. F1000Prime Rep 6:1–7. doi:10.12703/P6-52

Rainer J, Gatto L, Weichenberger CX. 2019. Ensembldb: An R package to create and use Ensembl-based annotation resources. Bioinformatics 35:3151–3153. doi:10.1093/bioinformatics/btz031

Rodnina M V. 2016. The ribosome in action: Tuning of translational efficiency and protein folding. Protein Sci 25:1390–1406. doi:10.1002/pro.2950

Sanchez M, Lin Y, Yang CC, McQuary P, Rosa Campos A, Aza Blanc P, Wolf DA. 2019. Cross Talk between eIF2α and eEF2 Phosphorylation Pathways Optimizes Translational Arrest in Response to Oxidative Stress. iScience 20:466–480. doi:10.1016/j.isci.2019.09.031

Saxton R, Sabatini D. 2017. mTOR Signaling in Growth, Metabolism and Disease. Cell 168:960–976. doi:10.1016/j.cell.2017.02.004.mTOR

Schuller AP, Green R. 2019. Roadblocks and resolutions in eukaryotic translation. Nat Rev Mol Cell Biol 19: 526–541. doi:10.1038/s41580-018-0011-4

Senju S, Iyama K-i., Kudo H, Aizawa S, Nishimura Y. 2000. Immunocytochemical Analyses and Targeted Gene Disruption of GTPBP1. Mol Cell Biol 20:6195–6200. doi:10.1128/MCB.20.17.6195-6200.2000

Shoemaker CJ, Green R. 2011. Kinetic analysis reveals the ordered coupling of translation termination and ribosome recycling in yeast. Proc Natl Acad Sci U S A 108:E1392–8. doi:10.1073/pnas.1113956108

Showkat M, Beigh MA, Andrabi KI. 2014. mTOR Signaling in Protein Translation Regulation: Implications in Cancer Genesis and Therapeutic Interventions. Mol Biol Int 2014:1–14. doi:10.1155/2014/686984

Spencer PS, Siller E, Anderson JF, Barral JM. 2012. Silent substitutions predictably alter translation elongation rates and protein folding efficiencies. J Mol Biol 422:328–335. doi:10.1016/j.jmb.2012.06.010

Stein KC, Frydman J. 2019. The stop-and-go traffic regulating protein biogenesis: How translation kinetics controls proteostasis. J Biol Chem 294:2076–2084. doi:10.1074/jbc.REV118.002814

Sudmant PH, Lee H, Dominguez D, Heiman M, Burge CB. 2018. Widespread Accumulation of Ribosome-Associated Isolated 3’ UTRs in Neuronal Cell Populations of the Aging Brain. Cell Rep 25:2447–2456.e4. doi:10.1016/j.celrep.2018.10.094

Thommen M, Holtkamp W, Rodnina M V. 2017. Co-translational Protein Folding: Progress and Methods. Curr Opin Struct Biol 42:83–89. doi:10.1016/j.sbi.2016.11.020

Thoreen CC, Chantranupong L, Keys HR, Wang T, Gray NS, Sabatini DM. 2012. A unifying model for mTORC1-mediated regulation of mRNA translation. Nature 485:109–113. doi:10.1038/nature11083.A

Wolf AS, Grayhack EJ. 2015. Asc1, Homolog of Human RACK1, Prevents Frameshifting in Yeast by Ribosomes Stalled at CGA Codon Repeats. RNA 21:935–945. doi:10.1261/rna.049080.114.frameshift

Woo K, Kim T, Lee K, Kim D, Kim S, Lee H, Kang H, Chung SJ, Senju S, Nishimura Y, Kim K. 2011. Modulation of exosome-mediated mRNA turnover by interaction of GTP-binding protein 1 (GTPBP1) with its target mRNAs. FASEB J 25:2757–2769. doi:10.1096/fj.10-178715

Wortel IMN, Meer LT van der, Kilberg MS, Leeuwen FN van. 2017. Surviving Stress: Modulation of ATF4-Mediated Stress Responses in Normal and Malignant Cells. Trends Endocrinol Metab 28:794–806. doi:10.1016/j.tem.2017.07.003

Yamashita R, Suzuki Y, Takeuchi N, Wakaguri H, Ueda T, Sugano S, Nakai K. 2008. Comprehensive detection of human terminal oligo-pyrimidine (TOP) genes and analysis of their characteristics. Nucleic Acids Res 36:3707–3715. doi:10.1093/nar/gkn248

Yu CH, Dang Y, Zhou Z, Wu C, Zhao F, Sachs MS, Liu Y. 2015. Codon Usage Influences the Local Rate of Translation Elongation to Regulate Co-translational Protein Folding. Mol Cell 59:744–754. doi:10.1016/j.molcel.2015.07.018

Zhang P, McGrath BC, Reinert J, Olsen DS, Lei L, Gill S, Wek SA, Vattem KM, Wek RC, Kimball SR, Jefferson LS, Cavener DR. 2002. The GCN2 eIF2 Kinase Is Required for Adaptation to Amino Acid Deprivation in Mice. Mol Cell Biol 22:6681–6688. doi:10.1128/mcb.22.19.6681-6688.2002

Zinoviev A, Goyal A, Jindal S, Lacava J, Komar AA, Rodnina M V., Hellen CUT, Pestova T V. 2018. Functions of unconventional mammalian translational GTPases GTPBP1 and GTPBP2. Genes Dev 32:1226–1241. doi:10.1101/gad.314724.118

Zinzalla V, Stracka D, Oppliger W, Hall MN. 2011. Activation of mTORC2 by association with the ribosome. Cell 144:757–768. doi:10.1016/j.cell.2011.02.014

